# Genuinely Blind Identification of Sleep Spindles through Trispectral Modulation Analysis

**DOI:** 10.64898/2026.08.24.746784

**Authors:** Christopher K. Kovach, Stephen V. Gliske, Leslie C. West, Junjie Liu, Michael O. Summers, Sukhbinder Kumar, Julie Gonzales, Owen Cox, Eric W. Tsang, John A. Thompson, Clete A. Kushida, Aviva Abosch

## Abstract

Sleep spindles, transient 11–16 Hz oscillatory bursts, are a defining electrographic feature of non-rem (NREM) sleep and a key biomarker of sleep physiology. A need for efficient and reliable identification of spindles motivates a large literature on automated detection algorithms. In this literature, annotation by trained sleep specialists remains the gold standard against which automated methods are trained, tuned and evaluated. However, inter-scorer agreement among experts is modest, which leaves a significant role for subjective judgment in the definition of a spindle. Finding objective, scorer-independent, criteria for identifying spindles remains an unresolved challenge. We report here a robust, highly specific, and previously unrecognized signature of spindle activity in the fourth-order spectrum (trispectrum), from which we identify the presence of spindles, characterize their waveforms, and obtain an optimal detection filter through a decomposition of the trispectrum (HOSD). Although it is a strictly blind, data-driven method, HOSD-based spindle identification and detection agrees well with expert annotation (median AUROC ~ 0.9), yet identifies many more events at the native threshold than both human scorers and comparison detectors. Many of these additional detections are confirmed as meeting AASM spindle criteria by four blinded specialists, demonstrating that spindle-like oscillatory bursting is prevalent below conventional human and automated detection thresholds. We observe that N2 sleep is distinguished principally by high-amplitude bursts, while low-amplitude bursting persists throughout NREM sleep, being globally suppressed only in REM sleep. We also describe robust identification of recording-specific spindle waveform properties such as frequency deceleration.

**Key Messages:**

- We introduce the first purely data-driven machine-learning algorithm for identifying and detecting sleep spindles that is independent of human judgment.
- This method identifies the presence of oscillatory bursting in the spindle band using information in the fourth-order spectrum (trispectrum) through a spectral decomposition of excess kurtosis in the spindle band.
- Detection thresholds are chosen based on a principled statistical criterion.
- This new method detects many more spindles than human scorers and other automated spindle detection algorithms.
- We present evidence that a large proportion of the additional detections represent true positives rather than false positives.
- We conclude that N2 sleep is primarily characterized by high-amplitude spindles, but that oscillatory bursts within the sigma band are prevalent below the conventional detection threshold in other non-REM stages and wakefulness, and uniquely suppressed during REM sleep.

## Introduction

Since their description by Loomis and colleagues in the 1930s [1], sleep spindles, prominent oscillatory bursts of 11-16 Hz that are essential for identifying and scoring N2 sleep, have emerged as a singularly informative biomarker of sleep physiology and pathology [2; 3; 4; 5; 6; 7; 8; 9]. Understanding their neural and behavioral correlates depends on reliable detection. The need for efficient and objective procedures has produced a large literature on automated spindle detection; yet, within this literature, visual scoring and review by a registered polysomnographic (sleep) technologist and a board-certified sleep medicine physician remains the standard against which automated methods are tuned and evaluated. However, agreement among expert scorers and between scorers and algorithmic detectors is typically modest [10], creating uncertainty about the true prevalence of spindles as well as appropriate detection thresholds for both humans and algorithms [11; 12; 13]. A crucial obstacle to progress on this question has been the absence of any objective criterion for identifying spindles that is not dependent on human judgment, either indirectly through human-labeled training data or directly through visual annotation.

Automated spindle detectors can be tuned to agree with the consensus of expert scorers at a level that approaches that of an individual expert [14; 15; 16; 13], yet it is not obvious whether such a consensus gives a more accurate gold standard or merely a more conservative one. Methods that seek to emulate human scorers must inherit biases and unstated assumptions of the human scorers. Among these is the, often implicit, adoption of an amplitude threshold, which is not specified among standard American Academy of Sleep Medicine (AASM) criteria for spindle identification in the scoring of N2 sleep [17]. In this context, the demonstration of a specific measure of spindle-related oscillatory bursting and a fully unsupervised method for isolating a signal on the basis of that measure would carry significant implications both for both the study of spindles and sleep staging, as variability in the assignment of electrographic features reduces inter-scorer reliability in sleep staging.

All existing spindle detectors make prior assumptions about the target signal with varying restrictiveness, forming a scale along which they might be arranged. At one extreme are methods that adopt a fixed pre-specified filter and threshold [18; 19; 20], while those that adjust to the statistical structure of the data extend along the scale toward the other extreme [21; 22; 23; 24; 25; 26; 27] (Table 1). Relevant parameters might be pre-calibrated against human consensus, informally through trial-and-error or through the formal application of supervised machine learning [28; 29; 30; 31; 32; 33; 34; 35], thus remaining, in all cases, tethered to human judgment. Measurement necessarily involves assumptions of one form or another, such as signal identifiability or a general range of frequencies in which the signal is contained. By “genuinely blind” we mean a method that makes no specific prior assumption about the target signal that does not also apply to a broad potential space of non-target components. It is explicitly the job of an unsupervised algorithm to separate one from the other without imposing assumptions within the shared signal space. Methods that rely on pre-filtering within the spindle band (e.g. [27]) are therefore regarded as implicitly supervised in the present classification, even if they do not involve formal training on labeled data.

**Table 1.** Sleep-spindle detection methods organized by signal isolation method (row), and parameter selection method (column). **Hard-coded a priori**: parameter values are fixed in advance. **Adaptive**: parameter values are tuned within a preset range according to specific prior assumptions about spindle characteristics (e.g. frequency of 11-16 Hz). **Trained on labels**: parameter values are tuned by supervised learning against expert spindle annotations. **Fully data-driven**: parameter values are determined from the data, with neither pre-specified thresholds nor human labels.

| Signal isolation method | Hard-coded a priori | Adaptive (strong priors) | Trained on human labels | Fully data-driven<br>(minimal priors, no labels) |
| --- | --- | --- | --- | --- |
| Bandpass + amplitude criterion | [18; 36; 37; 19; 38; 21; 39; 20; 40] | [24; 41; 42] | [43; 44; 29; 13; 45] | — |
| T-F decomp. (atomic or multitaper) | [22; 23; 25] | [46; 47; 48] | — | — |
| Learned feature representation | — | [27] | [30; 28; 31; 32; 33; 49; 35; 34; 50] | — |
| State-space oscillator model | — | [26] | — | — |
| Blind deconvolution based on HOS | — | — | — | **HOSD (this work)** |

The approach introduced here relies on estimation of a higher-order spectrum (HOS); specifically, a subdomain of the fourth-order spectrum, or *trispectrum*. It differs from prior methods in that both the presence and average characteristics of spindles in time and frequency emerge directly from higher-order spectral estimates without imposing prior assumptions about those characteristics. It therefore solves the problem of identification in a completely unsupervised manner. Because spindle characteristics are recovered from time-averaged statistics, rather than individual events, a detailed characterization of aggregate properties of the spindle waveform is obtained *before any spindles have been detected*. All previous methods for characterizing spindle waveforms begin with the detection of candidate events, for which specificity and biases of the initial detection inevitably affect the recovery of the target signal [12]. One consequence is that the presence of oscillatory bursts may be identified even under levels of noise that preclude the confident detection of individual events.

There is mounting evidence that fine-grained properties of spindles, such as frequency, duration, amplitude, scalp topography, and the internal modulation of each event, carry physiologically meaningful information about the underlying sources. Spindle density, duration, and peak frequency vary systematically with age and development [6], are altered in schizophrenia [19; 51; 8; 7], Parkinson’s disease [9], obstructive sleep apnea [52], and epileptic encephalopathies [35; 53], and they have shown promise as predictive biomarkers in early development [54]. Intra-spindle properties, such as frequency deceleration have emerged as potential biomarkers of the functional integrity of the thalamocortical loops that generate them [55; 56; 57; 58; 59; 60]. By recovering the signal waveform in a manner that is unsupervised, robust and independent of detection, the present method contributes a new toolkit for the accurate characterization spindle alteration under pathology.

## Methods

### Data

All data were obtained with permission from two open databases of polysomnography (PSG) recordings composed of intervals in which sleep spindles were independently annotated by one or more sleep specialists and one or more automated spindle detection algorithms.

#### MASS data

100 sets of full-night PSG data were obtained with permission from datasets S2-S5 of the Montreal Archive of Sleep Studies database (MASS)[16], accompanied by spindle annotations contained within the Massive Online Data Annotation (MODA) dataset, which included 3 to 10 intervals of 115 seconds in which sleep spindles were annotated according to each of the following criteria:[13] 1) a consensus of researchers, 2) a consensus of expert scorers, and 3) a consensus among a large sample of non-expert scorers recruited through crowd sourcing. The 100 recordings used in the present analysis correspond to the younger cohort (Phase 1) of the MODA spindle dataset, composed of healthy participants aged 18–35 years (mean 24.1) [13].

#### DREAMS data

The DREAMS Spindle database [61] includes PSG from 8 subjects with sleep spindles annotated within arbitrarily chosen 30-minute windows in a selected EEG channel (C3 or Cz) for each subject. Only data in the annotated window were included in the present analysis both for spindle estimation and performance calculation. In addition to the comparison detectors, comparisons included annotations provided with the database from two trained sleep experts and the output of an automated spindle detector tuned to match the human scorers’ performance [39].

#### Signal pre-processing

For each MASS recording, the channel scored by MODA scorers (C3, referenced to linked ear), was downsampled from 256 to 128 Hz, high-pass filtered at 0.1 Hz using a high-order FIR filter, and cleaned of large-amplitude transients by iteratively rejecting samples beyond six standard deviations from the thresholded sample mean until none exceeded the threshold. The DREAMS analyses likewise used each excerpt’s annotated channel (C3 or Cz, referenced to A1) with the same steps.

### Use of Higher-Order Spectra in Signal Identification

In the same way that the variance of a time series (average power) can be decomposed by frequency to yield the power spectral density (PSD), higher cumulants, like skewness and kurtosis, admit Fourier-domain decompositions as so-called higher-order spectra (HOS). As with the PSD and ordinary autocorrelation, HOS are Fourier-domain representations of higher-order autocorrelations, arising from the expectation of the product of the time series with itself at multiple time lags [62]. The *K*^th^-order HOS is a function of *K* − 1 frequencies; the spectral decomposition of kurtosis, which yields the fourth-order spectrum, or trispectrum, is therefore a function of three frequencies. Unlike the PSD, HOS preserve recoverable information about underlying waveforms encoded in Fourier-domain phase. The higher dimensionality of HOS also facilitates signal identification and separation: whereas PSD estimates contain *O*(*N/*2) coefficients for an *N*-sample signal, HOS contain *O*(*N* ^*K*−1^*/*(2*K*!)) coefficients, meaning that they are over-complete (containing more coefficients than the number of samples in the input signal) [63], with the practical implication that HOS estimates admit a PCA-like additive decomposition (HOSD) in a way that the PSD does not [64].

As cumulants, HOS are additive for an additive superposition of independent processes, thus an additive decomposition like HOSD can be interpreted as a model of independent component processes that generate the observed signal. Because higher-order cumulants vanish for stationary Gaussian processes, they are especially useful for recovering non-Gaussian signals from Gaussian noise, so that the HOS estimates represent the signal HOS without noise-related bias. Although neural sources are generally non-Gaussian, recordings of neural sources at any given location and moment in time often reflect a foreground of one or a small number of prominent sources embedded in a background that is a summation over a large number of distant or weak sources with similar amplitude, which, by the central limit theorem, tends towards a Gaussian distribution [65; 66].

#### Identifying Modulated Oscillations with the Trispectrum

Higher-order spectra, like the trispectrum, tend to be less open to an intuitive grasp than the power spectrum, but some insight might be developed by understanding the trispectrum as a measure of the linear association of time-varying spectral power across frequency bands (that is, the cross-spectrum of power between different frequency bands in the signal). In this formulation, two dimensions arise from the respective frequency bands and the third from the frequency of association between them [67; 68]. Information in the phase of the trispectrum encodes dynamic properties of the association; for example, making it possible to distinguish frequency from amplitude modulation [69], and in general to recover characteristic temporal patterns of modulation.

The high dimensionality of the trispectrum also places constraints on computational feasibility, while the representation and interpretation of a three-dimensional function remains inherently challenging. These difficulties may be met by limiting estimates to relevant lower-dimensional subdomains within the trispectrum. Oscillatory bursting in EEG represents the transient modulation of a carrier oscillation; its presence in a time series can therefore, in principle, be identified from the trispectrum, provided that the characteristics of modulation are sufficiently stable over time to permit reliable estimation of the relevant statistics. Modulation of a narrowband carrier oscillation by a real-valued time-varying envelope is revealed by correlated sidebands on either side of the carrier band, originating in the symmetric negative and positive halves of the spectrum of the envelope, recentered at the carrier frequency [70]. Modulation therefore leaves a signature in the statistical association between sidebands, which can be used to identify the presence and spectrotemporal characteristics of a modulated oscillation. In the context of a modulated carrier oscillation whose envelope is non-negative or lowpass, such that energy is preserved in the carrier band, relevant information is concentrated around a two-dimensional subdomain of the trispectrum representing the three-way association of carrier and side bands, which can be approached in isolation. A representation of this subdomain, the *modulogram*, reveals the spectrum of the carrier and the envelope along orthogonal axes in a directly interpretable form [68]. Finally, the relationship between the trispectrum and kurtosis implies that a modulated Gaussian carrier may be filtered in such a way that the output will have excess static kurtosis (i.e. kurtosis greater than 3) for any time-varying envelope. In the output of such a filter, the timing of modulation is revealed in large excursions that contribute to excess kurtosis. A natural threshold for detecting the presence of the target signal is obtained against a null model which assumes only Gaussian noise, that is, the threshold at which excess kurtosis vanishes in the residual.

### Higher-order spectral decomposition

A further step toward the practical recovery and interpretation of information in the trispectrum is obtained through an additive decomposition (HOSD), which explains the observed trispectrum in terms of a series of feature waveforms [64]. HOSD is motivated by a model of the signal as a mixture of independent linear processes driven by stationary super-Gaussian white noise (Lévy process), *u*_*i*_(*t*), conceived here for simplicity as a marked Poisson process where amplitude is the mark space. Each process is associated with a distinct finite waveform, *f*_*i*_(*t*), in a background of additive Gaussian noise, *n*(*t*):

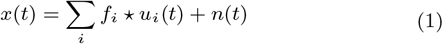

The *K*^th^-order autocorrelation of each component processes is given by

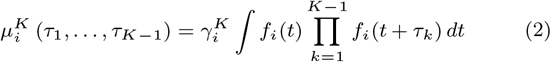

and the *K*^th^-order moment spectrum is obtained from a Fourier transform of *µ*^*K*^ :

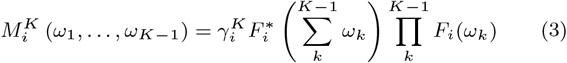

where *F*_*i*_ is the Fourier transform of *f*_*i*_, and *γ*^*K*^ is the moment of the driving process.

Cumulant spectra have the same form as (3) except within regions of infinitesimal support where subsets of frequencies sum to 0; in practice, these regions may be excluded from estimation with asymptotically negligible loss of information. Within the cumulant domain of *M* ^*K*^, two important properties hold: (1) HOS of the Gaussian noise process vanishes and (2) the separate spectra of the mixture components are additive:

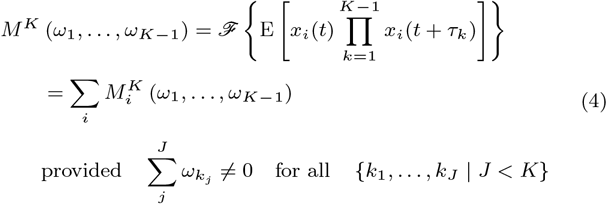

These properties motivate the additive decomposition of HOS as a method to separate independent component processes from each other and from additive Gaussian noise, under a rationale very similar to that by employed by independent component analysis (ICA) [71]. In practice, such a decomposition will tend to merge processes that emit very similar waveforms into a single component, while any individual process that exhibits significant variability of its emitted waveform will tend to be split between multiple components. For this reason, it is often useful to regard HOSD as recovering a feature space spanning the output of underlying processes. HOSD is, however, robust to violations of several assumptions expressed in Eq. (1): 1) for a colored driving process, *u*, that varies slowly or with infrequent changes of state relative to the scale of the estimation window (e.g. an inhomogeneous point process with slowly or intermittently changing rate), the autocorrelation structure of *u* is concentrated near the non-cumulant domains of HOS excluded in the estimate, which minimizes any influence it might have on the result. 2) For a colored driving processes with autocorrelation at the scale of the estimation window (e.g. a periodic point process), the autocorrelation structure will tend to be transferred to the feature waveform (e.g. *f* becomes a periodic train of *f* ‘s), which does not invalidate the model. 3) If a process is non-stationary such that *f* varies over time, the estimated form of *f* will represent an average if the variation is small, or conversely, if the variation is large, the estimate will be split over multiple components. In the former case, HOSD recovers a central tendency of the process; in the latter, HOSD recovers a feature space spanning different states of the process.

### Implementation of HOSD

Implementation of HOSD starts with the estimation of HOS normalized such that magnitude falls in the range [0, 1], which represents the strength of cross-frequency coupling at the given combination of frequencies. The direct method of HOS estimation [62] parallels Welch’s method for PSD estimation by segmenting a signal into a series of overlapping windows, tapering each window to mitigate spectral leakage, and averaging HOS computed on each segment over segments. Although several normalization schemes are applicable in the transformation to tricoherence [72], one particularly straightforward option quantifies the consistency of the HOS phase spectrum over segments (i.e. phase locking) by normalizing against the mean magnitude spectrum, representing the magnitude of the estimate that would be observed if phase were constant across all samples [73; 74]. As mentioned, the coupling among four frequencies quantified by tricoherence can be reframed as coupling between two pairs of frequencies, which in turn relates to power-power coherency across frequency bands in the Wigner-Ville time-frequency decomposition [68]. Tricoherence encodes both strength of coupling, in the tricoherence magnitude spectrum, and temporal characteristics, in the phase spectrum. As a measure of the statistical strength of coupling, tricoherence implicitly conveys signal-to-noise ratio in a way that is useful for obtaining an ideal matched filter [64], which accounts both for the phase spectrum of the target signal and its amplitude response relative to the noise background [75].

#### Matched filter and feature waveform estimation

HOSD for spindle detection was implemented in the following steps:

1. Segment the data into a series of 10 s analysis windows and taper to suppress spectral leakage.
2. Reject segments containing artifact or extreme outliers.
3. Compute the fast Fourier transform (FFT) of each segment.
4. Estimate the sample trispectrum as an average of segment trispectra. Estimates were obtained only for coefficients within the diagonal slice and specified target frequency ranges (5 Hz to 20 Hz carrier *×* 0.2 Hz to 3 Hz modulation frequency).
5. Normalize each coefficient by the average segment magnitude trispectrum to obtain tricoherence.
6. Project an outer product of each segment’s FFT onto the second and third dimensions of the sample tricoherence estimate (restricted to the diagonal slice) to obtain a partial filter estimate for each segment.
7. Average the segment filter estimates over segments to obtain an initial matched filter estimate.
8. Filter each segment with the matched filter estimate.
9. Circularly shift segment filter estimates to align the greatest extrema in the filter output across segments.
10. Multiply each segment filter estimate by the sign of the segment extremum.
11. Update the filter estimate as the average over realigned and sign-corrected segment filter estimates.
12. Repeat steps 7 through 12 until convergence through a maximum of 25 iterations.

This process converged rapidly on a filter matched to a component feature waveform within the additive mixture, with amplitude response weighted to signal-to-noise ratio (SNR) [64]. The feature waveform itself was obtained by averaging over filter-aligned and sign-corrected segments of the unfiltered data.

#### Component signal estimation

Under the super-gaussian linear model of (1), the component signal was understood as a series of possibly superimposed feature waveforms with varying amplitude and sign. The component was estimated in the following steps:

1. The previously obtained matched filter was applied to the unsegmented signal.
2. A threshold on the filter output was chosen such that excess kurtosis vanished within sub-threshold samples. Sub-threshold samples were set to 0.
3. The component signal was obtained as the convolution of the thresholded filter output and feature waveform estimate.
4. The result was scaled to minimize mean-square error between the filtered data and filtered component estimate.

Subsequent components were estimated through deflation, by repeating matched filter and component estimation on the residual signal after subtracting the component signal from the raw data for the second component, or the previous residual for all subsequent components.

### Event detection and segmentation

Event detection began with the thresholded filter output obtained in step 2 of component estimation, in which each oscillatory burst is typically associated with multiple threshold crossings. To obtain a detection signal whose peaks reflect the timescale of each oscillatory burst (rather than a single oscillatory period), root-mean-square power was computed as a moving average on the thresholded filter output using a kernel that matched the characteristic timescale of the filtered component waveform. This scale was estimated by applying the matched filter to the estimated feature waveform, which resulted in a symmetric zero-phase oscillatory impulse whose width conveys the timescale of the peak generated by a single detection. To suppress residual baseline noise in the feature impulse, the impulse was thresholded according to the kurtosis thresholding criterion and clipped to a symmetric window centered on the peak containing 99% of the total remaining energy. The timescale was then estimated as the standard width of the clipped and thresholded impulse. A Hann window smoothing taper with standard width matched to that of the impulse, was applied to the square of the thresholded filter output for the original signal; peak detection was then applied to identify event times. To minimize ringing in the smoothed signal, which might result in extraneous peaks, the smoothing taper was chosen to be compact, unimodal and continuously differentiable, properties satisfied by the Hann window. Resulting peak time may no longer align precisely with extrema in the unsmoothed filter output, so peak event amplitude was obtained through an approximation to a max filter, implemented as a weighted *L*^10^ norm: the thresholded filter output was raised to the tenth power, smoothed with the previously obtained Hann window, and the tenth root of the result at the event time was taken as peak amplitude. Event amplitude was normalized to the standard deviation of the sub-threshold filter output to give an estimated signal-to-noise ratio (SNR) for each event.

### Per-sample posterior probability estimation

In the previously described point-process model, the signal is treated as a series of events representing instances of the feature waveform in a background of noise. Given a model of the signal and noise distributions and an estimate of the event rate, the posterior probability that any given sample comes from a signal-containing interval may be computed. Estimating the respective model parameters must account for the uncertainty of signal and noise segmentation that motivates the estimate in the first place. This was done through an iterative procedure, expectation maximization (EM), which alternated between maximum-likelihood estimation of the respective distribution parameters with each sample weighted according to the probability of its membership in the respective class, followed by re-estimation of sample probability weights according to the updated mixture parameter estimates. For the present application, noise and signal amplitude distributions are both modeled as log-normal. The parameters of the noise distribution were estimated under a weighting of samples according to the local density of samples below the HOSD kurtosis threshold (presumed to represent noise). Noise parameters were subsequently held fixed throughout the EM iteration which updated only signal distribution parameters and the signal prior, *π*. The choice of a log-normal model for both noise and signal was arrived at by comparing a goodness-of-fit measure, Bayesian Information Criterion (BIC), between several alternative specifications, which included combinations of gamma and log-normal for noise amplitudes as well as a non-parametric (histogram-based) signal distribution. BIC favored the log-normal model in the majority of MASS recordings (median ΔBIC ≈ −4 *×* 10^5^).

The per-sample posterior *P*_*t*_(signal|*z*_*t*_) was obtained from the mixture model fit to the smoothed detection signal, *z*_*t*_. Each sample is drawn from a signal containing interval, with prior probability *π*, or from non-signal (noise only) interval, with probability 1 − *π*,

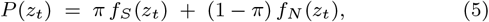

where *f*_*N*_ and *f*_*S*_ are both log-normal densities of the form

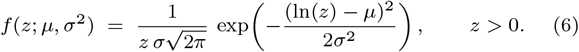

The noise parameters (*µ*_*N*_, *σ*_*N*_) are obtained from a weighted maximum-likelihood estimate (MLE) on the unthresholded smoothed-RMS distribution and held fixed throughout EM iteration; only the signal parameters (*µ*_*S*_, *σ*_*S*_) and the prior *π* are subsequently updated. Each sample *z*_*t*_ receives an initial noise responsibility weight based on the fractional contribution of sub-threshold samples to local power as obtained with the previously described moving-average kernel:

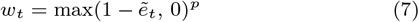

where 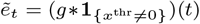 is the convolution of the supra-threshold indicator with the smoothing kernel *g* (i.e., the fractional contribution of samples above the kurtosis threshold to local power), so that 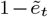 is the corresponding sub-threshold fraction. The oscillatory nature of the signal leaves a substantial fraction of sub-threshold samples within signal-containing intervals, around the zero crossings of *x*^thr^, regardless of SNR; an ad-hoc remedy that sharpens the distinction between signal and noise intervals was applied through exponent *p*, set to *p* = 3.

Estimation of the signal distribution under the EM algorithm alternated the following steps:

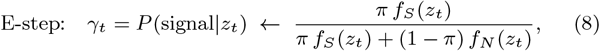

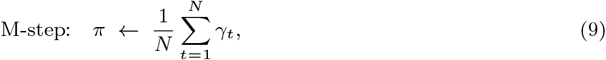

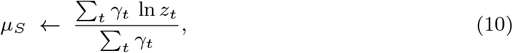

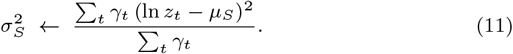

To reduce the computational burden of direct estimation over a large number of samples, EM was not applied to the raw input data, but to binned counts within 500 log-spaced bins over the range of *z*_*t*_, with *P* (signal|*z*_*t*_) evaluated at each bin center. Binned parameter estimates typically diverged by less than 0.1% from those obtained from un-binned data while reducing computing time for whole-night recordings by two orders of magnitude. EM parameter estimation discarded samples with *z*_*t*_ *<* 0.1 to suppress artificial spikes near 0 created by artifact rejection and other sources of data dropout. The M step estimated signal distribution parameters, weighting data according to the signal distribution responsibility, *γ*_*t*_, as obtained in (5). The E step updated responsibility weighting according to the updated parameter estimates. E and M steps alternated until the relative change in log-likelihood fell below 10^−6^. The algorithm was initialized by setting signal responsibility to the initial weighting derived from the cumulant threshold in (7), 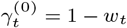.

The fitted mixture model yielded the sample-level posterior probability *P* (signal | *z*_*t*_) estimate. For each detected event the posterior at the event’s peak time was taken as an estimated probability that the event represents a true positive: 100 detections each with *p* = 0.2 should yield about 20 true positives in expectation, so weighting events by *p* produces a model-implied true-event count.

#### Spindle segmentation

We compared two approaches to segmenting spindles from background based on the output of the HOSD detector, one based on peak detection in the HOSD filter output, thresholded at the cumulant threshold, and the second based on the output of the mixture model described in the previous section.

##### Peak-based segmentation

The first approach, (HOSD_peak_) applied simple peak detection to the cumulant-thresholded filter output, with event onset and offsets selected at the full-width-at-half-maximum (FWHM) crossings on either side of the peak, as described in *Event detection and segmentation*, with additional merging of adjacent peaks. A candidate spindle epoch was identified as the extent of the smoothed detection signal around each peak that is greater than half the peak amplitude. Candidates whose half-maximum windows overlapped were merged into a single event. Merged events took as their extent the union of half-maximum windows, and as magnitude, the amplitude of the greatest peak. The peak-based approach is maximally permissive, typically introducing many events with comparatively low signal-to-noise ratio (SNR*<* 3) and correspondingly low estimated posterior probability.

##### Posterior-probability thresholding

In the second approach, segmentation relied directly on the sample-wise posterior probability estimate, *γ*_*t*_, that a given sample fell within a spindle interval as described in *Per-sample posterior probability estimation*, with event onsets and offsets determined by threshold crossing rather than peak detection. The samplewise posterior was first smoothed with a 0.25 s moving average, then thresholded at a set probability threshold, *θ*. Event onset and offsets were identified according to threshold crossings, and events of duration less than 0.3 s or greater than 3 s were discarded. Performance of the probability-based HOSD detector was evaluated at two different operating points: HOSD_p85_ uses *θ* = 0.85, which was near *F*_1_-optimal operating point against the MODA expert consensus (peak pooled *F*_1_ = 0.642). The second detector, HOSD_p50_, adopts the Bayesian maximum a posteriori (MAP) threshold, *θ* = 0.50, which yields a high recall / low precision detector against the MODA expert consensus.

### Statistical validation

Inferential tests on trispectral coefficients within the modulogram were derived from mass univariate tests on individual coefficient estimates, obtained by average HOS over multiple segments of the signal. The relevant test is on the mean of the complex-valued coefficient, for which a complex-valued general linear model (GLM) was applied [76]. To relax parametric assumptions of the GLM, cluster-based procedures [77] were applied to correct for multiple comparisons: the test statistic within clusters of coefficients exceeding a pre-specified significance threshold in the mass-univariate parametric tests were summed, and a surrogate null distribution of cluster statistics was constructed from the maximum cluster values across surrogate estimates. Clusters were defined as groups of contiguous coefficients with parametric uncorrected significance below the *α* = 0.05 threshold, and summed within-cluster deviance served as the test statistic, which was referred to a null distribution of 1000 samples from the phase-randomized surrogate distribution. By randomizing phase but preserving the magnitudes of sample HOS, this procedure retains bias arising from the distribution of magnitude in the surrogate distribution [74], as well as smoothing bias in the estimator.

#### Cumulative analysis

To examine the SNR threshold at which events are reliably present, we computed cumulative cluster statistics over analysis windows sorted by estimated signal magnitude. Data segments used in computing HOS estimates were sorted in ascending order according to the maximum SNR magnitude within the detection filter output for the segment. Cumulative statistics were then computed on sorted segments. For this analysis, the sigma-band-associated cluster boundary (shown in Fig. 7A) was set according to the threshold at which 90% of datasets exhibited a significant difference from 0 at false discovery rate (FDR) corrected *Q <* 0.05. The test statistic, computed per dataset, was the sum of within-cluster deviances for the univariate complex-valued general linear model (GLM). A cumulative null distribution was obtained through phase randomization: the per-subject 95^th^ percentile of the *N* = 200 permuted null cumulative means defines a per-subject null threshold for the observed mean. A cohort threshold curve was obtained as the median across subjects of the per-subject null curves, against which the observed cohort-median curve was compared at each sorted SNR value.

### Performance evaluation

Both DREAMS and MASS databases were used to evaluate the performance of the three HOSD-based detectors (HOSD_peak_, HOSD_p50_, HOSD_p85_) against comparison detectors.

#### Comparison detectors

Comparison detectors applied to both databases were those of Ferrarelli et al. [19], Kramer et al.[35], the a7 detector of Lacourse et al. [29], Martin et al. [40], and OpenSpindleNet of Sejak et al. [50]. For the DREAMS excerpts these were supplemented by the automated annotations of Devuyst et al. provided with the database [39], DETOKS of Parekh et al. [25], and Spindler of Larocco et al. [47]; for the MASS recordings, by the detectors of Mölle et al. [36], Wamsley et al. [20], and He et al. [26]. DETOKS and Spindler were restricted to the DREAMS excerpts: DETOKS performed poorly on the MASS recordings at preset parameter values, and would have required additional tuning of parameters to perform adequately, while the quadratic runtime of Spindler made the application to all 100 whole-night recordings impractical. In both cases, we opted not to apply *ad hoc* adjustments that would have been necessary to include them in the analysis. Annotations by the human scorers provided within each database were included as additional responders.

#### Event clustering and consensus boundaries

Following [29], for both DREAMS excerpts and MASS samples, events were matched across detectors and annotations according to pairwise overlap at a threshold of 20% (intersection-over-union ratio, IoU *>* 0.2). To determine the event boundaries for the gold-standard-free latent-class analysis of MASS data and to specify candidate events in the DREAMS validation analysis, candidate events were identified within the pairwise coincidence matrix as clusters (connected subgraphs) involving one or more detections across participating detectors. For each cluster, the event onset and termination were taken as the median of the respective values reported by participating detectors. Events reported by each detection were then matched to the consensus event in which they participated. This procedure means that it is possible for a detection within a given cluster to not overlap with the consensus interval at an intercept-over-union (IoU) threshold of 0.2; such discrepancies were, however, infrequent, affecting fewer than 5% of detections.

#### Event-based performance evaluation

For the MASS database, performance metrics were computed in two ways: (1) The Massive Online Data Annotation (MODA) expert scorer gold standard (GS) computed performance against the consensus among experts provided in the MODA platform [13] and (2) A GS-free latent class model (LCM) which derived a consensus among multiple detectors / responders by fitting a model in which the true class was treated as an unobserved latent variable. For the DREAMS excerpts, performance analysis was based on *post-hoc* expert scorer evaluation of candidate events, which were either consensus detections among available detectors or foil events which did not overlap with any individual detection.

##### MODA expert scorer GS

For precision-recall and receiver operating characteristic (ROC) analyses on the MASS data, HOSD candidate events were matched directly to gold-standard events following [29], using IoU matching with an IoU threshold of 0.2. When multiple events from a given detector or responder matched a given GS event, the event with greatest IoU was assigned as the matching event and remaining events treated as unmatched. Likewise, when multiple GS events matched a single detection, only the event with the greatest IoU ratio was counted as a match. Matched pairs are true positives (TP), unmatched events are false positives (FP), and unmatched GS events are false negatives (FN). True negatives (TN) were sampled by pairing each GS event with a randomly selected interval of the same duration that did not overlap with any GS event. For the posterior probability detector HOSD_pp_, matching was based on detection cluster membership. For the peak-based HOSD detector, precision-recall curves were obtained by sweeping a threshold over estimated event SNR and recomputing TP, FP, FN and TN at each threshold.

##### GS-free analysis of MASS data

For the GS-free LCM (Fig. 4, see *Latent class model*), event clusters served as the observation units, with a detector or responder’s event matched to any consensus event in which it participates. Consensus event class was assigned as true if the expectation of the latent state exceeded 0.5 for that event. Although performance may be estimated directly from the LCM parameter fits, for the sake of consistency with the expert GS, reported performance metrics were based on an LCM consensus GS (*ŷ*_*j*_ ≥ 0.5 using the IoU *>* 0.2 matching criterion.

##### DREAMS validation

For the DREAMS validation (Fig. 6, see *Performance validation with the DREAMS data*), the evaluator’s binary judgment (rating *>* 0.5) provided the truth label for both consensus events and foils. For each combination of evaluator, detector and excerpt, a subset of consensus and foil events were classified as positive or negative according to the evaluator’s judgement. Unrated events were excluded from the analysis.

In each case, true/false positives/negatives were tabulated and converted to performance metrics:

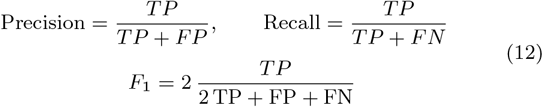

#### Sampling of negative cases

The event-based matching schemes above yield true positives, false positives, and false negatives, but provide no true negatives, and therefore cannot support estimation of specificity, false-positive rate, or area under the receiver-operating characteristic curve (AUROC). To recover a true-negative class, we constructed a matched negative pool for each subject in the MASS data. The empirical distribution of that subject’s GS positive events was sampled with replacement to produce *N* negative events, with *N* equal to the positive count (positive-to-negative ratio 1:1). Each negative event was placed at a uniformly random start time within the annotated PSG segments under two constraints: (i) the sampled event does not overlap any GS positive event, and (ii) consecutive sampled events are separated by at least 0.5 s. Positive events and sampled negative events in the GS set were counted as true (respectively false) positives if a detected event overlapped it with IoU *>* 0.2 and otherwise as a false (respectively true) negative. From the resulting (score, label) pairs we computed per-subject ROC curves using the sampled negative cases to compute TNR and FPR, ignoring detections that did not coincide with an event in the true class or randomly sampled negative class. Because precision and recall do not depend on TNR, precision–recall curves and the area under them (AUPRC) were computed using all detections, with every detection not matching a GS positive counted as a false positive. Chance AUPRC thus equals each GS’s positive base rate.

#### Linear mixed-effects modeling for DREAMs data

For each performance metric, a linear mixed-effects model (LME) was fit across all scorers, detectors and excerpts, which included a random intercept for excerpt.

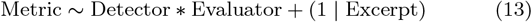

Analysis was conducted in MATLAB R2023b (`fitlme`). Because the number of scorers was smaller than the size recommended for grouping variables in LME models [78], random effects for evaluator were not included in the model, but the influence of evaluator was included as a fixed effect. Per-detector marginal means were obtained as contrasts over the fixed effects. Pairwise contrasts of HOSD detection against every other detector were tested via `coefTest` and adjusted within metric with the Holm–Bonferroni procedure.

### Latent class model

Because experts exhibit substantial disagreement in the spindle detection, expert rating provides an imperfect GS. We therefore complemented the expert GS with a GS-free latent-class model (LCM), which treats the true class as a latent variable. The LCM interleaves estimation of the sensitivity and bias of each responder with the estimation of the true class of each event, where each responder’s verdict is weighted according to the current estimate of its performance [79; 80]. In the absence of a GS event demarcation, candidate events were defined as clusters of detections among the available detectors. Clusters were identified through graph-based clustering as connected subgraphs within the graph whose edges represented pairwise overlapping detections among all pairs of detectors. Pairwise overlap was defined as before according to IoU *>* 0.2 (see *Event clustering and consensus boundaries*).

In the LCM model, each candidate is assumed to either contain a spindle or not (*y*_*j*_ ∈ *{*0, 1*}*), and the output of each responder, *r*_*ij*_ is a conditionally independent detection:

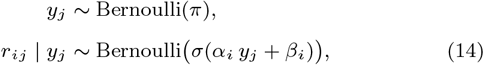

where *σ*(*u*) = (1 + *e*^−*u*^)^−1^. The bias parameter, *β*_*i*_, is the log-odds false positive rate (FPR):

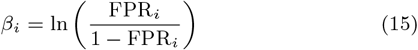

the discrimination parameter *α*_*i*_, is the log-odds ratio of true positive rate (TPR) to FPR, which quantifies how informative the responder is:

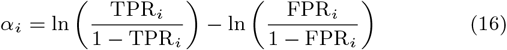

Moderate *α*_*i*_ with strongly negative *β*_*i*_ describes a detector with high specificity but low sensitivity, while high *α*_*i*_ and moderate *β*_*i*_ describes a detector with high sensitivity but low specificity. A detector with *α*_*i*_ = 0 is entirely uninformative, being equally likely to generate a detection for true and false events.

The LCM model was fit using 11 responders on MASS data: the eight algorithmic detectors (HOSD_p85_, Ferrarelli, Martin, Mölle, Wamsley, Lacourse (A7), Kramer, He) and the three MODA scorer subgroup consensuses (expert, researcher, non-expert). HOSD_p85_ is the posterior-probability variant of HOSD with events thresholded at *γ*_*t*_ ≥ 0.85, which approximates the MODA expert GS threshold. Annotators not included in the LCM fit, including individual MODA scorers, HOSD_peak_, and HOSD_p50_ (the same posterior-probability variant thresholded at *γ*_*t*_ ≥ 0.5), were scored against the latent consensus at a threshold of Pr(*y*_*j*_ = 1 | *r*_*·j*_) ≥ 0.5.

The LCM model was fit by EM [79] with alternating expectation (E) and maximization (M) steps. The E step provides a responsibility weight (*ŷ*_*j*_ = E [*y*_*j*_ |*r*_.*j*_]) given the output of all responders weighted by the current estimate. The informativeness of each responder was estimated in the M step under the current responsibility weighting given by the preceding E step. The E-step computed *ŷ*_*j*_ = E [*y*_*j*_] from the current (*α*_*i*_, *β*_*i*_, *π*) according to the sum of log-likelihood-ratios across observed responders:

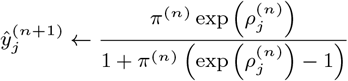

where

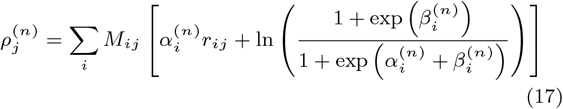

where *M*_*ij*_ ∈ *{*0, 1*}* represents a mask that limits the estimate to the evaluated window among responders that do not apply to the full data set (e.g. MODA scorers).

The M-step updated each responder’s (*α*_*i*_, *β*_*i*_) according to the regularised weighted-mean TPR and FPR estimates:

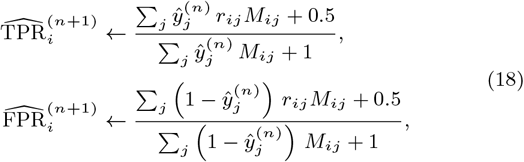

where the +0.5/+1 regularizer acts as a prior on TPR and FPR that prevents parameters from blowing up in data sets with few events or detections. The latent base rate updated as *π*^(*n*+1)^ ← mean(*ŷ*^(*n*)^). Iteration continued until max_*j*_ |Δ*ŷ*_*j*_ | *<* 10^−7^ (typically *<* 50 iterations). Ambiguity of sign in the parameters is resolved by initializing *ŷ*_*j*_ at the majority vote across responders.

#### SNR stratified analysis of the MASS data

Because HOS estimates average over repeated observation, they may be used to identify signals that do not manifest with sufficient SNR to be distinguished from noise in the raw data. To test for the presence of such “occult” spindles, which fall below detectability using standard methods, we applied trispectral modulation analysis to data stratified by SNR amplitude. This analysis focused on the subset of modulogram coefficients for which at least 90% of MASS samples reached FDR-corrected significance of Q *<* 0.05 in the whole-night recording. For each MASS sample, we computed the per-segment fourth-order spectrum at 77 trispectrum coefficients meeting this criterion, all of which fell within the sigma band. Segments were sorted in ascending order according to the maximum peak smoothed root-mean-square (RMS) of the un-thresholded component filter output. For each cumulative rank *k*, the statistical magnitude of the deviance of the mean coefficient was computed under a parametric complex-Gaussian model [76], and the sum of deviances over the 77 coefficients within the mask served as the figure of merit for determining the presence and magnitude of a statistical effect. As a negative control, the analysis was repeated at a region of the trispectrum centered on carrier frequency, 10 Hz, and envelope-modulation frequency, 2 Hz, where no FDR-significant coefficients were observed in ≥ 90% of samples. To obtain a threshold demarking the transition between detectable and undetectable “occult” spindles we computed the amplitude at which the signal-plus-noise mixture model assigned responsibility (posterior probability) greater than 0.5 to the signal process, as described in *Per-sample posterior probability estimation*. This represents the threshold at which a Bayesian detector identifies the presence of a spindle with ≥ 50% confidence.

#### Performance validation with the DREAMS data

We considered whether the sensitivity of visual annotation increases if scorers are asked to judge previously identified candidate events rather than detect and judge events by eye in raw data. To this end, a subset of candidate events identified by HOSD and comparison detectors were presented to each of four scorers along with non-spindle foils. To minimize the possible effect of a positive response bias, and to remain faithful to the imbalance of positive and negative cases within the raw data (intervals containing no spindle far exceed intervals that contain a spindle), most presented events (66% – 75%) were foils. For each detector, evaluator and excerpt, performance metrics, *F* 1, precision and recall were computed using the evaluator’s verdict as the GS.

The pool of candidate events used in this analysis was drawn from detections by HOSD, Devuyst, Martin, Ferrarelli, and the two DREAMS visual annotations, along with a set of foils placed at peaks in the 11–16 Hz envelope of that fell at least 2 s from any detected or scored spindle, each assigned a duration sampled from the HOSD detections. The remaining detectors (OpenSpindleNet, Kramer, Lacourse (A7), Spindler, and DETOKS) were added to the comparison after evaluators had completed their evaluations and so were scored against a fixed pool to which they did not contribute detections. This may raise a concern that the analysis is biased against the excluded detectors; however, it was observed that for all excluded detectors but one, detections that did not coincide with detections already in the set (i.e. discovered by at least one of the included detectors/scorers or chosen as a foil) constituted less than 2% of the total. The exception was the more permissive detector of Kramer et al., for which 30% of its detections did not coincide with events in the pre-existing set. Because only events reviewed by an evaluator were included in the analysis, the direction of any bias in performance estimates for the Kramer detector resulting from this exclusion depends on whether the excluded events would have been rated as spindles at a higher or lower rate than included events. A reasonable upper bound on precision in the excluded sample in this case is provided by the baseline precision among the rated Kramer detections (one may safely assume that events discovered by no other method are at least not more likely to be endorsed as spindles than events also discovered by one or more of the included detectors), while the baseline endorsement rate among rated foil events provides a reasonable lower bound on precision (events discovered only by Kramer and no others are, at a minimum, not less likely to be endorsed than those discovered by none of the included detectors). The range of biases in F1 suggested by these bounds is −0.054 to +0.055, within which the overall rank of Kramer with respect to the other detectors changes only near the extremes and only by one (from position 3 up to 2 or down to 4). We have therefore elected to include the additional detectors for the sake of completeness while retaining the original set of ratings.

## Results

### Spindle identification in the DREAMS database

We applied trispectral modulation analysis to all 8 segments of EEG data with spindle annotation provided in the DREAMS Spindle database [61]. Each recording consisted of a 30-minute PSG excerpt from a single frontal channel (C3 or Cz, referenced to A1) with spindles annotated by two expert scorers and by an automated detector [39]. Tricoherence was computed within the diagonal slice of the trispectrum over the carrier range 5–20 Hz and envelope range 0.2–3 Hz, and HOSD was applied to the resulting modulogram in each subject (Fig. 1).

**Figure 1.**
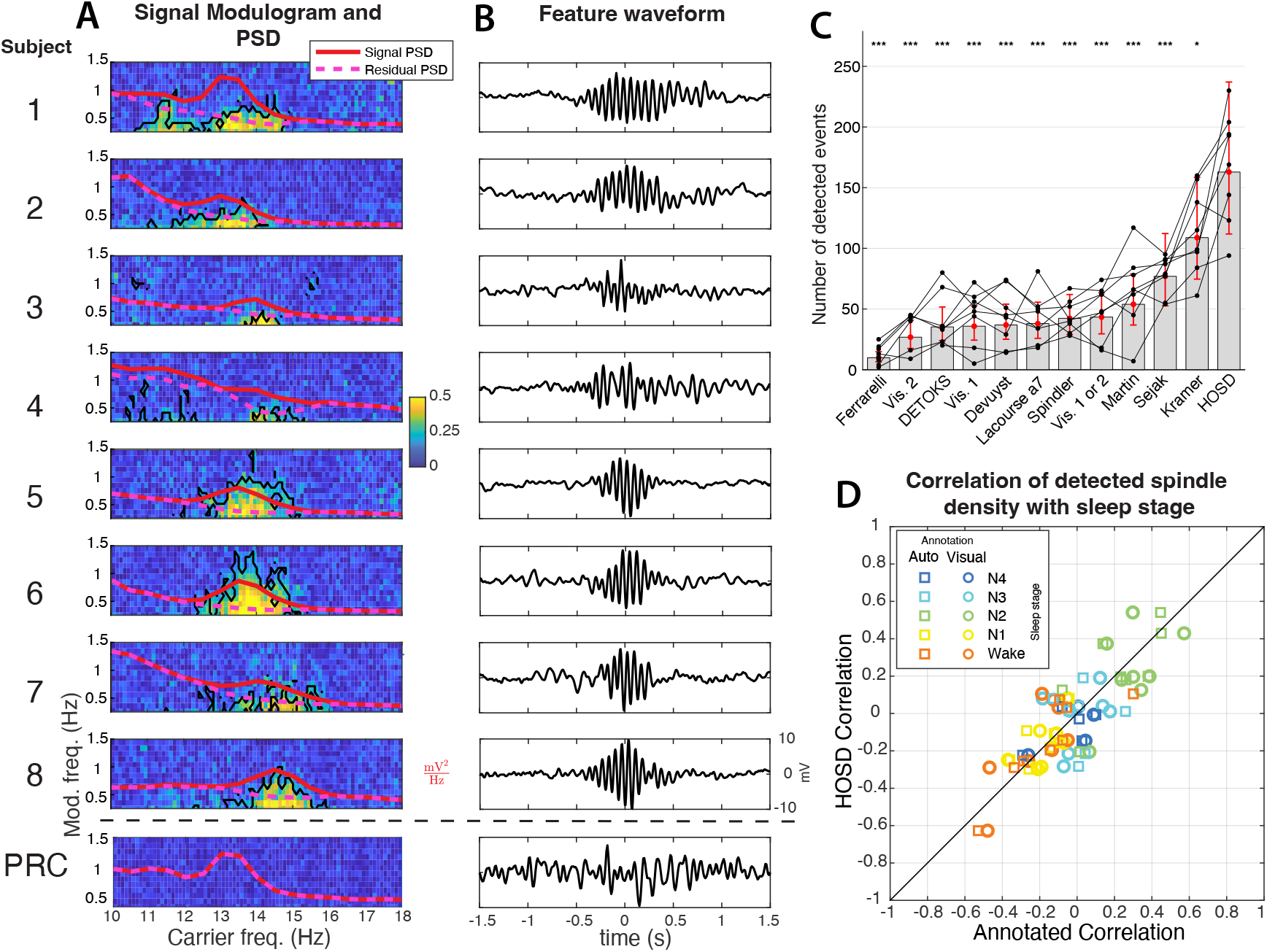
Comparison of blind sleep spindle identification with HOSD, visual scoring and an ad-hoc automatic algorithm. Eight samples of 30 minutes of polysomnography data were obtained from the DREAMS open database of polysomnography data [61] along with annotated sleep spindle times obtained through visual and an automatic detection algorithm [39]. Spindles were annotated in selected 30 min recording excerpts from a single frontal channel near C3 or CZ, referenced to A1. **A:** HOSD was applied to the modulogram estimated within the 5–20 Hz carrier range and 0.2 to 3 Hz modulation range on the same excerpt. For each subject modulograms (**A**, *left column*) exhibited one or more peaks in the sigma band (11-16 Hz) associated with sleep spindles (**A**, *black lines* indicate *P <* 0.05, cluster permutation test). Power spectral density (PSD) plots (**A**, *red*), likewise exhibited a peak in corresponding regions of the sigma band. In each case the first component returned by HOSD (**B**) showed spindle-like morphology with frequency and temporal scale of modulation corresponding to peaks in the associated modulogram. PSD plots of the residual signal, after removing the first HOSD component, show complete suppression of the peak in the sigma band (**A**,*magenta dashed lines*), indicating that it is explained by modulated power as identified in the modulagram. For comparison, the bottom plots in (**A** and **B**) show the result of applying HOSD to a signal in which temporal structure is removed through phase randomization (phase-randomized comparison, PRC). The sigma-band peak is preserved in the PSD but eliminated from the modulogram (**A**, *bottom panel*). HOSD returns a feature with irregular temporal structure (**B**, *bottom panel*), consistent with random noise. **C:** HOSD identifies a larger number of spindle events than both visual and automated annotation methods (Kruskal-Wallis *P <* 0.001). **D:** Comparison of the correlation between spindle density as estimated with HOSD (*y-axis*) and each annotation method (*x-axis*, square: automatic, circle: visual); correlations did not differ significantly between methods for any sleep stage (Kruskal-Wallis *P >* 0.5).

In every subject, the modulogram (Fig. 1A, *left column*) exhibited one or more peaks at carrier frequencies within the sigma band (11– 16 Hz) that were significant against a cluster-based permutation null (*P <* 0.05). Sigma band activity was also observed as a peak in the power spectral density (Fig. 1A, *red*), which coincided with the peak carrier frequency identified in the modulogram. The first component returned by HOSD (Fig. 1B) recovered, in each case, a waveform with spindle-like morphology whose carrier frequency and envelope timescale matched the location of the sigma-band peak in the associated modulogram. Removing this first component from the signal eliminated the sigma-band peak in the residual PSD (Fig. 1A, *magenta dashed lines*), indicating that the sigma-band power in these recordings is fully accounted for by component recovered through HOSD.

To illustrate that these results reflect temporal structure associated with modulation rather than stationary power, and to provide a null comparison, the same analysis was applied to a surrogate time series obtained by randomizing the Fourier domain phase of excerpt 1 (phase-randomized comparison, PRC). Phase randomization preserved the PSD of the surrogate, including the sigma-band peak, but eliminated the peak in the modulogram (Fig. 1A, *bottom panel*). HOSD returned a feature with irregular temporal structure, consistent with random noise (Fig. 1B, *bottom panel*), demonstrating that information conveyed by the modulogram is distinct from that conveyed by the power spectrum alone.

### Correlation with sleep stage

HOSD identified a greater total number of events compared to other detectors (Fig. 1C), with a median of 383.5 events per recording, compared with medians of 89 and 50 for visual and automated annotation, respectively (Kruskal–Wallis, *P <* 0.001). Despite this discrepancy in absolute counts, spindle density per sleep stage estimated by HOSD was strongly correlated with density estimated by each reference method, and these correlations did not differ significantly between the visual and automated references in any sleep stage (Fig. 1D; Kruskal–Wallis, *P >* 0.5). These results show that spindle-like events identified by HOSD exhibit the expected relationship to NREM sleep in the DREAMS database.

### Identification of spindles in the MASS database

To characterize trispectral properties of spindles more comprehensively in a large dataset and to evaluate whether observations from the DREAMS data generalize to prolonged recordings (whole night rather than 30 min.) covering multiple stages of sleep, we applied the same analysis to 100 whole-night recordings from the MASS database. Each recording in this dataset contains one or more brief (115 s) windows in which spindles are annotated by three groups of scorers, trained sleep specialists (exp), researchers (re), and a large cohort of non-expert scorers (ne). Modulograms were computed with data from channel C3 from each of 100 whole-night recordings from the MASS dataset. The complex-valued estimates were averaged across recordings to form a grand-average modulogram (Fig. 2). The magnitude of the average tricoherence (Fig. 2A) exhibited a single, well-circumscribed peak centered in the sigma band along the carrier axis, extending over a range of envelope frequencies consistent with the typical, 0.5 - 2 s, timescale of spindle modulation. Moreover, the sample average over recordings was highly representative of a pattern that was consistent across individual recordings: at a conservative false-discovery-rate threshold (*Q <* 0.05), each of the 100 recordings showed a tricoherence estimate significantly different from zero within the sigma band (Fig. 2B), indicating that sigma-band modulation is a reliable feature at the single-subject level. In addition to information about the statistical magnitude of modulation, the modulogram preserves information about the temporal characteristics of modulation at different frequencies by way of the phase modulogram (Fig. 2C), which exhibited a characteristic non-uniform distribution, again, highly consistent across datasets as shown by the magnitude of the average modulogram phase spectrum over datasets, with vector length near unity within the sigma band peak (Fig. 2D). Finally, modulogram estimate filtered by sleep stage showed a strong preference for N2 sleep, with smaller but significant peaks in N1, N3 and N4. Taken together, these results show that sleep spindles leave a stereotyped, statistically robust fingerprint in the trispectrum of spontaneous EEG.

**Figure 2.**
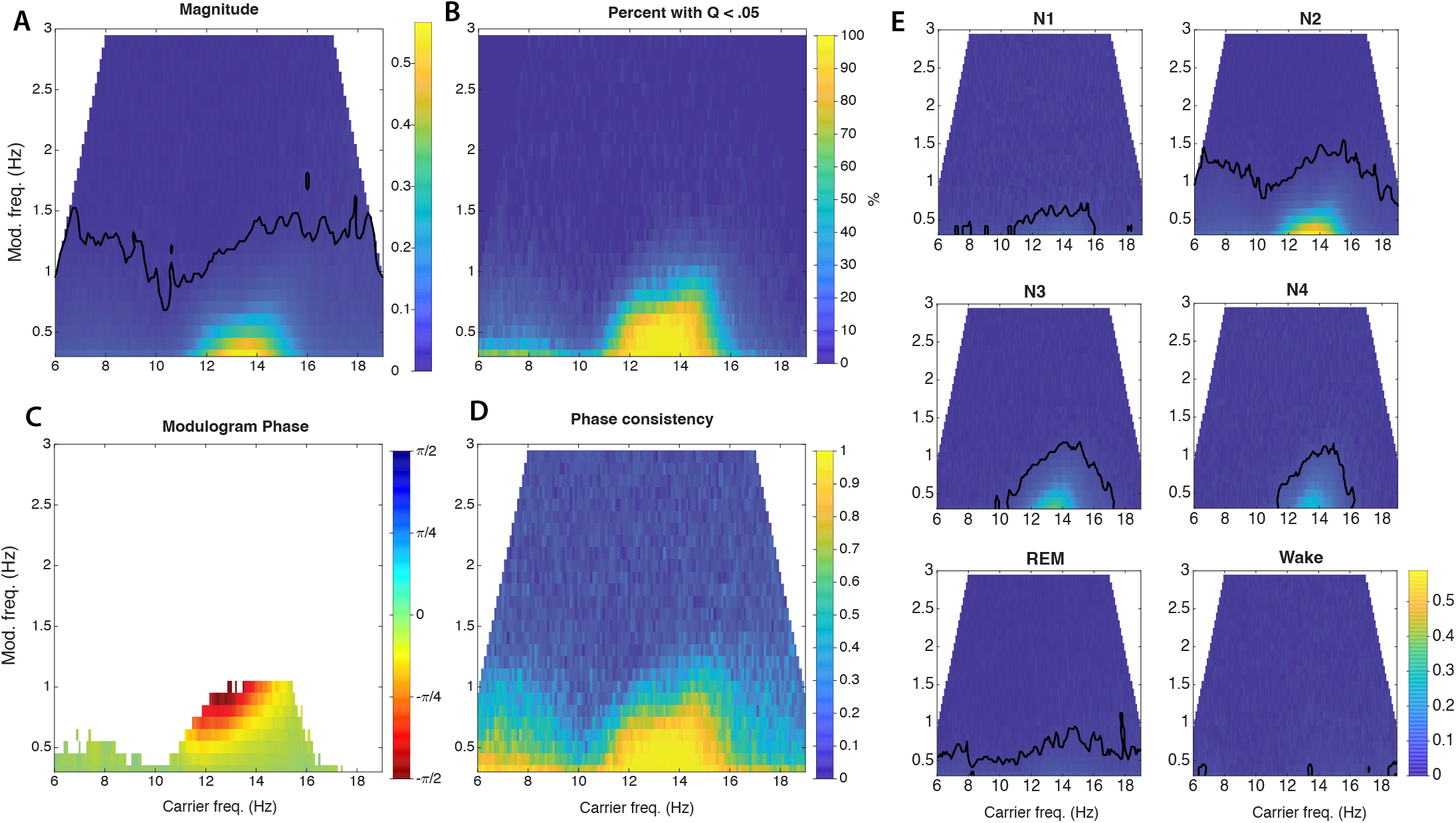
Spindle-associated modulation creates a prominent signature in the trispectral modulogram. Tricoherence was computed within the diagonal slice for each of 100 whole-night recording included in the MASS database. **A:** Magnitude of the average complex-valued tricoherence over all 100 samples. **B:** Percent of samples for which the whole-night tricoherence estimate differed significantly from zero at FDR threshold *Q <* 0.05. Every sample showed evidence of modulation within the sigma band at this conservative FDR corrected threshold. **C:** Phase angle of the average, thresholded within the region for which at least 25% of the estimates differed significantly from 0. A variable group delay across frequencies is revealed in the non-uniformity of phase, with relatively greater slope, hence greater delay, at low frequencies, corresponding to a characteristic frequency ramp (chirp) in the feature waveform (see Fig. 3A). **D:** Consistency of phase across estimates, measured as mean vector length of estimated tricoherence phase over recordings.

#### Spindle waveforms in the MASS dataset

To obtain a characteristic waveform associated with sigma-band modulation, HOSD was applied to each full-night recording in the MASS database. In every case, the first component recovered by HOSD was associated with the sigma-band peak in the modulogram, consistent with the result obtained with the DREAMS database. Individual waveforms obtained this way were subjected to a further round of HOSD to obtain an aggregate “grand average” waveform (Fig. 3B). The Wigner-Ville distribution of this grand-average feature (Fig. 3A) exhibited a negative frequency ramp, which is the time-domain counterpart of the carrier-dependent group delay seen in Fig. 2C, showing the latter as a correlate of intra-spindle deceleration. Intra-spindle deceleration was also highly consistent across individual recordings: a negative slope was observed in the waveform Wigner-Ville distribution (WVD) in 98 of 100 datasets (Fig. 3C) with a median, −3.9 Hz/s (IQR −4.6 to −3.0), significantly less than zero at the group level (Wilcoxon signed-rank test, *P* ≪ 0.001). In addition to intra-spindle deceleration, the grand average waveform revealed an (Fig. 3B) asymmetry of rise and fill times, with a shorter rise time than decay (Fig. 3D): measured as decay time minus rise time, this asymmetry was significantly greater than zero at the group level, median 0.38 s (IQR 0.14 to 0.78) (*P <* 0.001).

**Figure 3.**
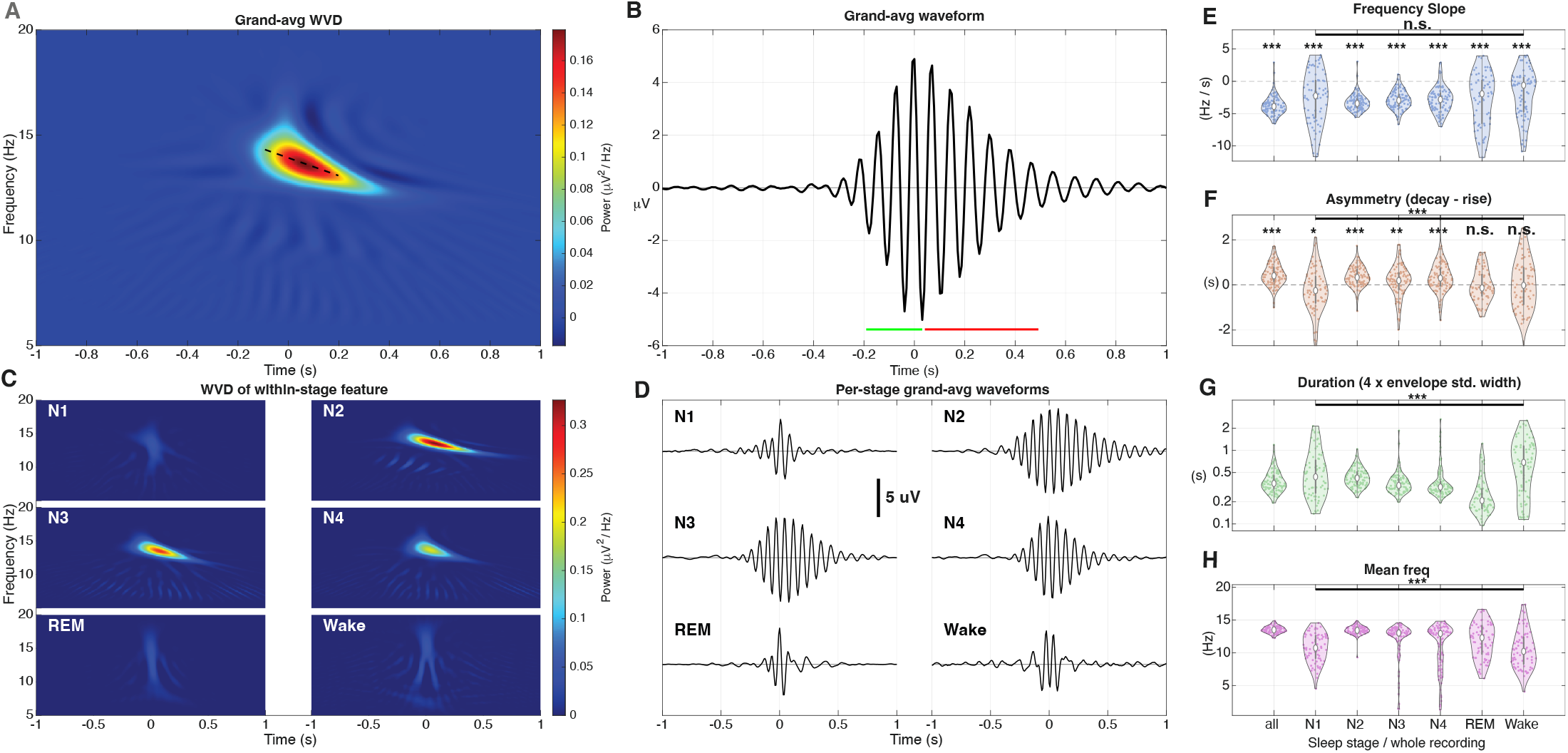
Grand-average feature waveform obtained by a second round of HOSD applied to 100 per-recording spindle waveforms from the MASS database. **A:** Wigner-Ville distribution (WVD) of the grand average feature waveform estimates over the whole recording. Variable group delay observed in Fig. 2C is reflected here in a negative frequency ramp (*dashed line*). **B:** Grand average feature waveform, demonstrating a characteristic asymmetry in the rise (*green line*) and decay (*red line*) phases. **C:** WVD plots of grand-average features estimated within respective sleep stages. **D:** Grand average waveforms over features estimated with respective stages. Grand average spindle waveforms were recovered from N2, N3 and N4. **E-H:** Violin plots showing the distribution across 100 spindle waveform estimates from the MASS database for frequency slope, asymmetry index (decay time minus rise time), duration as 4*×*standard envelope width, mean waveform frequency. Distributions are shown for waveforms estimated through HOSD over the whole recording, and for estimates including only the indicated sleep stages (*columns 2-7*). Bar and asterisks indicate significance of the main effect of sleep stag at *P <* .001(***), *P <* .01(**), *P <* 0.05(***), and *P >* 0.05 (n.s). Asterisks above violin plots [81] in **E** and **F** indicate significant difference of the mean from 0. **E:** A reliably negative slope was observed in waveforms estimated over the whole recording, and separately in stages N2, N3 and N4. Other stages exhibited variability but with significantly negative median (signed-rank test, *P <* 0.001) which did not differ across all stages (*P* = .08). **F:** Mean of asymmetry indices, measured as the difference between decay and rise times was positive (decay *>* rise) for the estimates over the whole recording, N2, N3 and N4 (*P <* .001). It was negative for N1 (*P <* 0.05) and non-significant (*P >* 0.05) in REM and Wake. **G:** Distribution of mean waveform duration exhibited significant variation over stage (*P <* .001). **H:** Distribution of mean waveform frequency exhibited significant variation over stage (*P <* .001).

#### Reliability of spindle identification

HOSD is not constrained to identify spindle-like features. To quantify whether identified features resembled spindles following AASM criteria [17], we assigned a “spindleness score” (S3), according to the degree of overlap with the spindle band (11 Hz – 16 Hz) and adherence to expected duration (0.25 s – 2 s). We observed a spindle-like oscillatory burst to be the dominant feature recovered by HOSD applied to each of the 100 whole-night recordings: spindleness scores fell within a range of [0.893, 1], with median 0.999, falling above 0.95 in 95% of cases. Note that, because the first component identified by HOSD exhibited a high spindleness score in every case, components were not sub-selected based on this score or any other spindle-related criterion. The prominence of spindle band modulation caused the algorithm to converge blindly on modulated oscillations within the spindle band in every examined dataset without further guidance. Thus we did not apply any additional tests to separate spindle from non-spindle features and conducted subsequent analyses on the first HOSD component recovered from each dataset. It is possible that additional components might recover spindle activity, e.g. separating high- and low-frequency spindles, but a detailed consideration of whether HOSD might separate spindle subtypes is left for future work.

#### Evaluation against the latent class

Because expert evaluation is an imperfect GS, we complemented MODA expert GS with a gold-standard-free latent-class model. The LCM was fit to detections from eleven responders (eight algorithmic detectors plus the consensus ratings of the expert, researcher, and non-expert MODA subgroups as given by Lacourse et al. 2020) in the MASS dataset. As described in *Latent class model*, the fitted model yields estimates of the sensitivity and bias of each responder along with the probability that each event represents a true instance of the latent class. The LCM consensus GS (used in Table 2 and Fig. 4C–D) was obtained by assigning events with *ŷ*_*j*_ *>* 0.5 to the “true” class.

**Figure 4.**
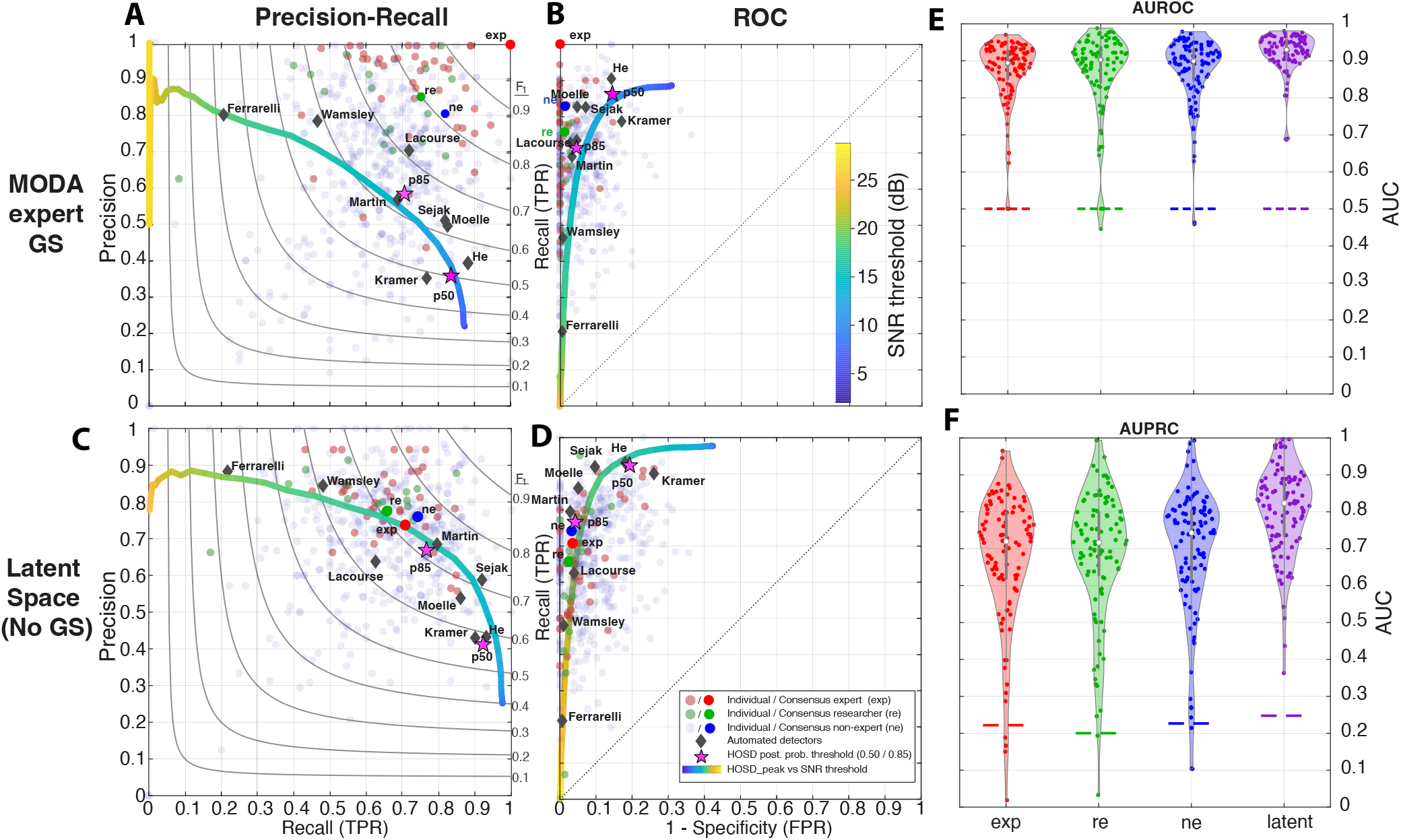
Performance against consensus GSs in the MASS database and latent space estimates. **A-D:** Precision-recall (**A**,**C**) and receiver operating characteristics (**B**,**D**) of automated detectors and individual MODA scorers, expert (exp), researcher (re), and non-expert (ne), against the MODA expert consensus (**A**,**B**) and a gold-standard-free latent space model (**C**,**D**). The latent-class model treats the true spindle state as an unobserved latent variable and estimates each responder’s sensitivity and bias together with the per-event posterior probability of a true spindle. Precision recall (PRC) and receiver operating (ROC) curves for peak based HOSD spindle identification are shown on respective panels with SNR amplitude threshold indicated by color scale. Estimates are obtained from all detections pooled across datasets. For precision (**A**,**C**) every detection that does not match a consensus spindle is counted as a false positive; the true negative set for the ROC (**B**,**D**), was obtained by randomly sampling a positive event and shifting it to a randomly sampled position with no event. Curved gray lines in PRC plot contours of the F1 statistic. **E-F:** Distribution of area under receiver operating (**E**) and precision recall (**F**) curves for HOSD_peak_ per recording estimated against respective GSs (latent: latent posterior probability thresholded at 0.5). Dashed lines mark chance: 0.5 for AUROC (**E**), and each GS’s positive base rate (≈ 0.20–0.25) for AUPRC (**F**).

**Table 2.** Latent class model fit for 12 responders used in fitting the model. Each row reports the fitted discrimination 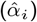 bias 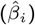, precision (*P*), recall (*R*), and (*F*_1_) of each responder against the LCM consensus (*ŷ*_*j*_ *>* 0.5). HOSD_p50_ was not included in fitting the LCM.

| Responder | $\hat{\alpha}$ | $\hat{\beta}$ | $P$ | $R$ | $F_1$ |
| --- | --- | --- | --- | --- | --- |
| $GS_{ne}$ | 4.59 | -3.43 | 0.76 | 0.74 | 0.75 |
| Martin | 4.21 | -2.63 | 0.68 | 0.80 | 0.74 |
| $GS_{exp}$ | 4.41 | -3.41 | 0.74 | 0.71 | 0.72 |
| Sejak | 4.22 | -1.51 | 0.59 | 0.92 | 0.72 |
| $HOSD_{p85}$ | 3.98 | -2.46 | 0.67 | 0.77 | **0.71** |
| $GS_{re}$ | 4.73 | -4.00 | 0.78 | 0.66 | 0.71 |
| Mölle | 4.45 | -1.65 | 0.54 | 0.86 | 0.66 |
| Lacourse (A7) | 3.21 | -2.52 | 0.64 | 0.63 | 0.63 |
| Wamsley | 7.77 | -7.74 | 0.84 | 0.48 | 0.61 |
| He | 3.22 | -0.20 | 0.43 | 0.93 | 0.59 |
| Kramer | 2.71 | +0.00 | 0.43 | 0.90 | 0.58 |
| $HOSD_{p50}^\dagger$ | 3.29 | -0.18 | 0.41 | 0.92 | **0.57** |
| Ferrarelli | 7.36 | -8.59 | 0.88 | 0.22 | 0.35 |

$^\dagger$ Held out from LCM fit.

Among the algorithmic detectors, Martin (*F*_1_ = 0.736), Sejak (*F*_1_ = 0.717), and HOSD_p85_ (*F*_1_ = 0.713) achieved the highest agreement with the latent consensus; the three MODA consensus detectors fell within a narrow band (*F*_1_ = 0.711–0.751, with the non-expert marginally higher than the rest), consistent with the finding that the three MODA subgroups are of comparable quality at the consensus level [13]. Although the remaining detectors exhibited lower F1 against the LCM consensus, comparison of Fig. 4C with the MODA expert GS in 4A shows a general shift of operating points towards a mutualy consistent precision-recall curve at points less balanced with respect to precision and recall (Fig. 4): Wamsley and Ferrarelli exhibited high precision and low recall, while Kramer and He achieved high recall at low precision. All detectors thus appeared to be tuned to the same latent class with comparable discrimination, differing mainly with respect to operating thresholding.

Several responders included in the performance analyses in Fig. 4 were held out of LCM fitting to avoid biasing the fit towards subclasses of potentially correlated detectors, such as variants of HOSD and individual members of the MODA scorer groups. Estimates for these responders were obtained through post-hoc comparisons to the LCM consensus estimate. HOSD_peak_ achieved AUPRC = 0.753 and peak *F*_1_ = 0.731 (at 15.5 dB SNR) against the latent consensus (Fig. 4C); the corresponding ROC, computed against a sample of spindle-free negatives matched to positives, gave AUROC = 0.940 (Fig. 4D). The HOSD_p50_ operating point (*P* = 0.411, *R* = 0.922, *F*_1_ = 0.568) fell near the range of high-recall low-precision detectors, near the maximum-a-posteriori (MAP) threshold.

### Spindle amplitude and sleep stage in the MASS database

To examine the relationship between the amplitude of HOSD-identified spindle events and sleep stage, events were stratified within thirteen 2-dB SNR bins between 5 and 29 dB. The density of events in each stratum was correlated with stage indicator variables separately for each MASS dataset. Fig. 5A shows the average *±*1 standard deviation over all 100 datasets. The relationship between N2 and spindle is reflected primarily in amplitude rather than density: spindles with amplitude in the range 15 – 21 dB were correlated with N2 sleep, *r* ≈ 0.35, whereas those below 15 dB, which constituted the bulk of detected spindle events (Fig. 5B) showed little selectivity for N2 over N3 and N4 sleep. REM correlation was negative at every amplitude, although the greatest suppression, *r* ≈ −0.35, appeared within the 13–18 dB range.

**Figure 5.**
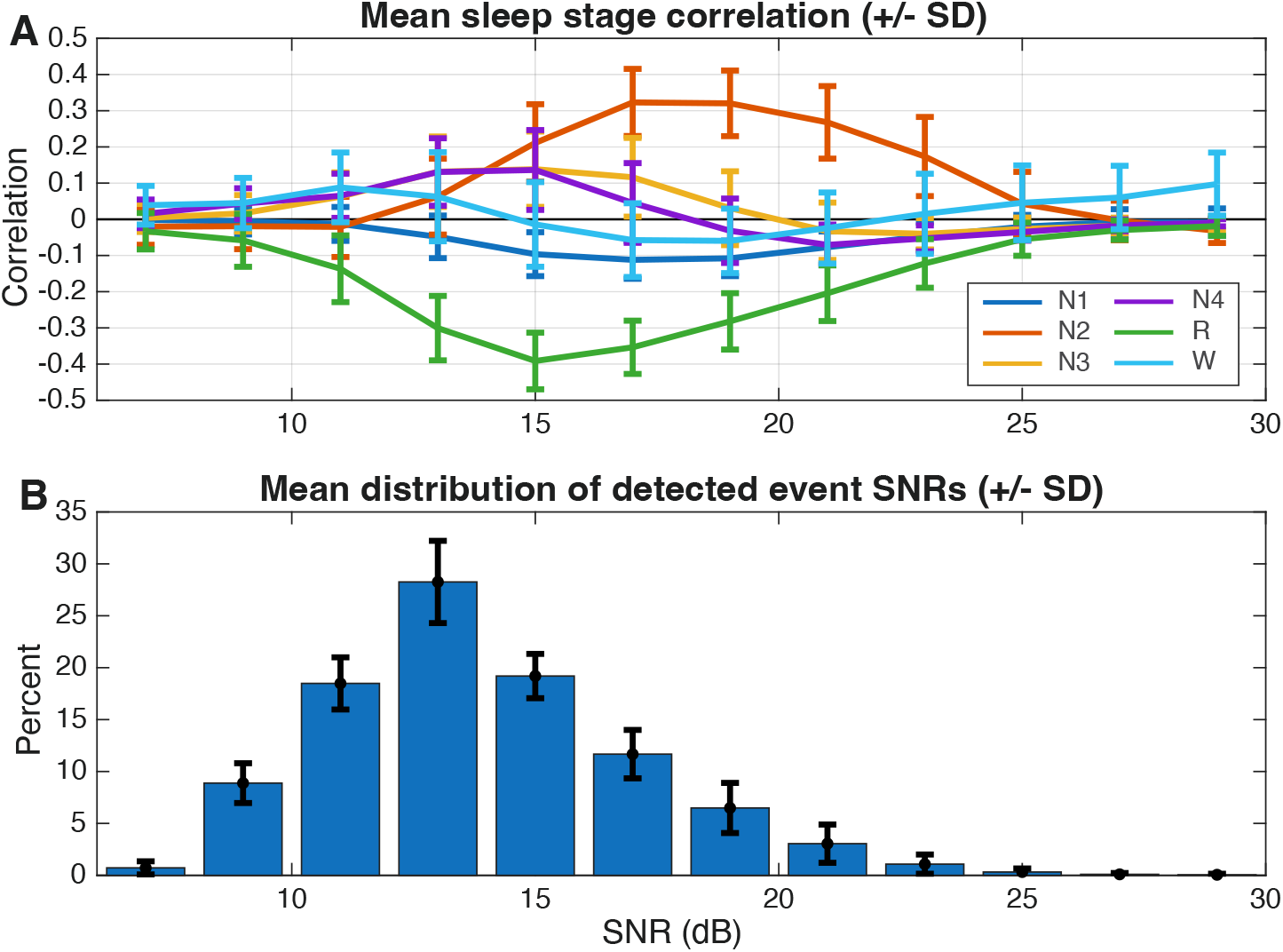
**A**: Correlation between spindle density and sleep stage across different relative amplitudes, broken down by SNR amplitude. Spindle counts for spindles falling within the given amplitude ranges were computed within each 30 s sleep-staging epoch, and correlated with sleep stage for each epoch, excluding unscored intervals. Lines show the mean correlation across 100 datasets and error bars show *±* 1 standard deviation of the data. Stage 2 NREM sleep is prominently associated with high-amplitude (*>*15 dB) spindles, while spindles of low and intermediate amplitude are non-specific with respect to NREM sleep. Suppression during REM sleep is evident at all amplitudes. **B**: Mean distribution of SNR spindle amplitudes *±* 1 std. dev.

### Validation with the DREAMS database

Whether events identified by a detector meet AASM criteria as sleep spindles is a distinct question from whether they match the performance of a human scorer. Performance comparisons against the human-scorer GSs inherently address the latter directly and the former only indirectly. For example, it remains possible that low-amplitude events that do not stand out obviously from background EEG are systematically overlooked by human scorers, even though they otherwise meet AASM criteria for sleep spindles (*which include no amplitude criterion*). To address this question, we recruited four expert sleep specialists to review a pool of candidate events pooled across detectors, along with “foil” events added to control for positive response bias. Each expert provided a rating on a scale from 0 (not a spindle) to 1 (definitely a spindle).

For the present analysis, ratings were binarized at 0.5 and perfomance metrics, recall, precision and F1, were computed against the binarized rating separately within each evaluator and 30 min. excerpt. These metrics were then compared across detectors using linear mixed effects modeling with evaluator as a fixed effect and random intercept grouped by excerpt. LME marginal means (Fig. 6A) place HOSD highest on *F*_1_ (0.55) and recall (0.71). OpenSpindleNet follows at *F*_1_ = 0.52, statistically indistinguishable from HOSD detection (Table 3), but at the opposite balance precision and recall: high precision (0.70) at markedly lower recall (0.46). Kramer is next at *F*_1_ = 0.48, then Martin (0.44) and Automatic detection (0.40). The remaining algorithmic and visual-scoring detectors cluster at *F*_1_ = 0.34–0.38; Ferrarelli is the lowest performer (*F*_1_ = 0.13) due to very low recall despite high precision. Detector main effects were highly significant for all three metrics (*p* ≪ 0.001). Holm-adjusted post-hoc contrasts of HOSD detection against every other detector were significant for *F*_1_ except for OpenSpindleNet and Kramer, whose contrasts did not survive Holm correction (Table 3). Although precision for HOSD detection was significantly lower than several detectors, its *F*_1_ score was dominated by much larger recall.

**Figure 6.**
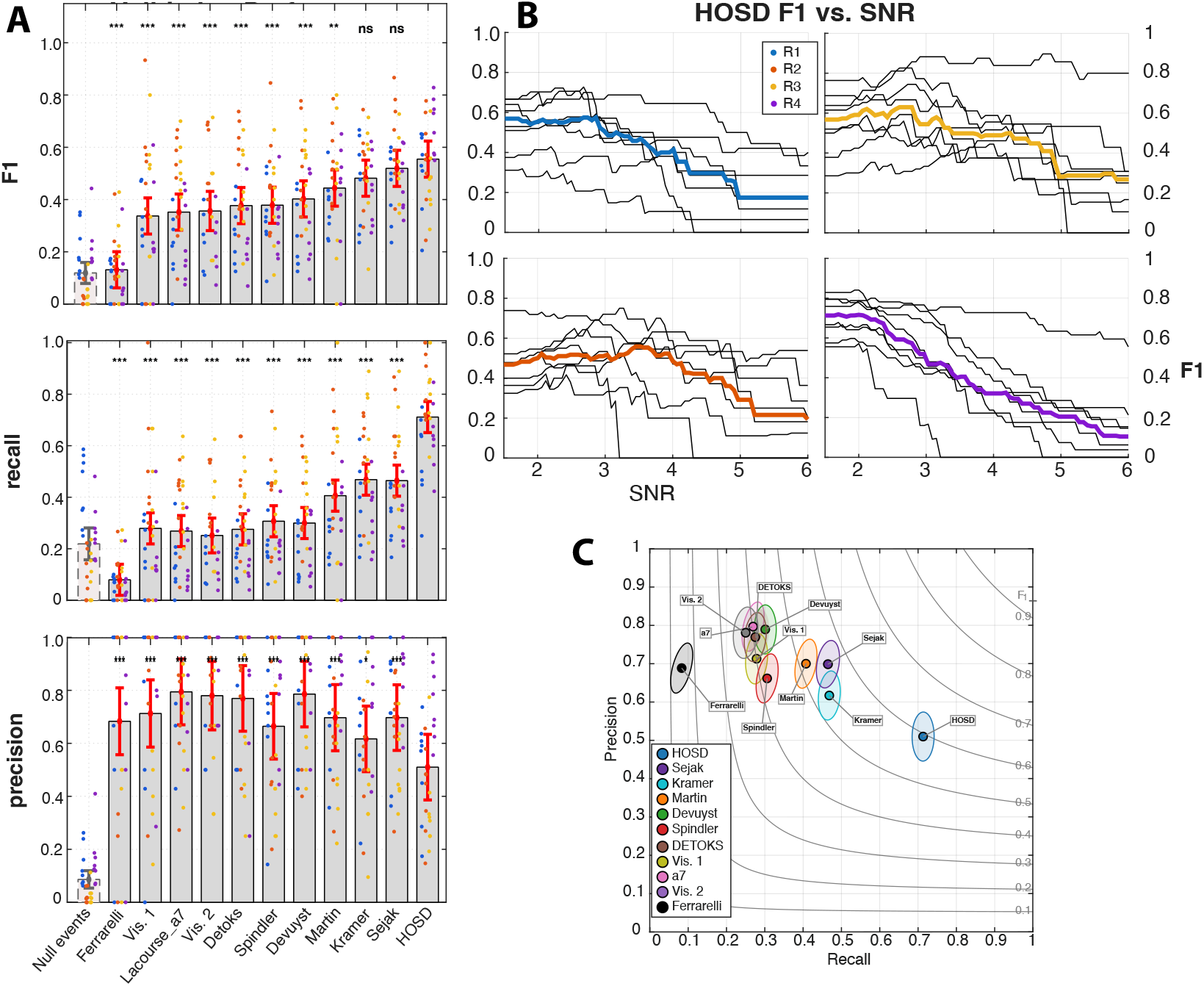
Expert validation of candidate spindles in the DREAMS database. Performance of HOSD_peak_ and alternative detectors was evaluated by presenting to each of 4 expert scorers (R1-R4) candidate spindles identified by any responder along with foils containing no detected events (null events). Scorersjudged whether or not the candidate event or foil represented a spindle. **A:** Dots represent performance metrics F1 (*top*), recall (*middle*), and precision (*bottom*) in each dataset against each evaluator’s judgment as the GS. Bars represent the fixed effect (*±* 2 SEM) of each detection obtained with a linear mixed-effects (LME) model that included fixed-effects for evaluator and detection method and random intercepts for dataset. Null events were not included in LME fitting; bars for the null events represent mean (*±* 2 SEM). Main effects of detection method and evaluator were significant (*P <* 0.001) with superior performance of HOSD against all other methods with respect to recall (*P <* 0.001) and all but OpenSpindleNet (Sejak) and Kramer with respect to F1 (*P <* 0.01). HOSD detection resulted in lower precision (*P <* 0.05) in 4 of of 7 comparisons, thus HOSD operated with higher recall but lower precision than alternative detectors in this analysis. **B:** F1 score vs. SNR threshold (linear scale) on HOSD detection against each evaluator’s judgment. For each evaluator, median performance (colored line) at the given threshold was computed across the 8 DREAMS datasets (black lines). **C:** Precision–recall operating point of each detector, showing LME estimates of mean recall and precision across evaluators and excerpts, with an ellipse indicating LME-derived dispersion, overlaid on iso-*F*_1_ contours.

**Table 3.** Contrast of HOSD marginal means from the LME against comparison detectors for *F* 1. Reported *p*-values are uncorrected; Holm-adjusted contrasts remain significant at *p <* 0.05 for every comparison except Sejak (2025) and Kramer (2021).

| Compared Detector | $\Delta$ | SE | $t_{(300)}$ | $p$ |
| --- | --- | --- | --- | --- |
| Sejak, 2025 | +0.035 | 0.034 | 1.03 | 0.31 |
| Kramer, 2021 | +0.073 | 0.034 | 2.12 | 0.035 |
| Martin, 2013 | +0.111 | 0.034 | 3.22 | $1.4 \times 10^{-3}$ |
| Devuyst, 2019 | +0.152 | 0.034 | 4.43 | $1.3 \times 10^{-5}$ |
| Spindler | +0.176 | 0.034 | 5.13 | $5.3 \times 10^{-7}$ |
| DETOKS | +0.178 | 0.034 | 5.17 | $4.3 \times 10^{-7}$ |
| Visual_scoring2 | +0.199 | 0.037 | 5.31 | $2.1 \times 10^{-7}$ |
| Lacourse_a7, 2018 | +0.203 | 0.034 | 5.90 | $1.0 \times 10^{-8}$ |
| Visual_scoring1 | +0.218 | 0.034 | 6.33 | $9.0 \times 10^{-10}$ |
| Ferrarelli, 2007 | +0.424 | 0.034 | 12.32 | $1.7 \times 10^{-28}$ |

The amplitude–*F*_1_ sweep (Fig. 6B) shows the expected inverted-U shape for evaluators R1, R2, and R3: *F*_1_ is roughly flat at low thresholds, peaks near 8–10 dB, then falls off as too many true spindles are removed. For R4 the threshold-zero *F*_1_ is already near maximum and the median curve decreases monotonically, indicating that R4’s positive ratings are well-explained by HOSD detection across its full amplitude range.

### Evidence for occult spindles in the MASS database

Conventional spindle detectors apply an amplitude threshold that has been calibrated, directly or indirectly, to expert visual annotation, with the implicit assumption that genuine spindles below this threshold are rare. Whether physiologically meaningful spindle activity persists below the visual-detection threshold (occult activity) is difficult to address because any method that relies on thresholding is inherently blind to subthreshold events. Trispectral modulation analysis overcomes this limitation because the underlying HOS estimators suppress noise through time-averaging, which allows for identifying signal characteristics in aggregate data even when individual instances of the signal appear at sufficiently low SNR to be undetectable.

To address the prevalence of occult spindles, we sorted segments used in HOS estimation by the maximum of the standardized HOSD detection filter output, expressed in dB, within the predefined population-level modulogram sigma cluster (Fig. 7**A**). We then evaluated the cumulative cluster sum-of-deviance from the intercept-only complex GLM over each ascending cumulative subset, comparing the observed curve at each sort-axis position to a null distribution derived from *N* = 200 phase-randomization permutations of the per-segment HOS coefficients (coefficients for each segment rotated by a random phase). This procedure yields a conservative estimate of the amplitude at which evidence of spindles begins to emerge.

Across all *n* = 100 MASS subjects, the sum-of-deviance measure crossed the *p* = 0.05 threshold of the null distribution in the vicinity of cumulant detection threshold, which in every case fell well below the Bayesian 0.5 posterior threshold used by HOSD_p50_ (Fig. 7**B**, red). The median sum-of-deviance curve crossed the median phase randomized null threshold at 8.66 dB, coinciding with the cumulant detection threshold (cohort median 8.64 dB, IQR 8.53– 8.72 dB). The median first-crossing at the subject level fell somewhat lower 7.75 dB (IQR 5.25–9.00 dB). The HOSD_p50_ window-crossing threshold lay ~ 3.8 dB above the cluster crossing (median 12.45 dB, IQR 12.08–12.80 dB). The out-of-band comparison cluster (carrier 10 Hz, modulation 2 Hz; Fig. 7**B**, black) tracked the null threshold curve throughout, never accumulating cluster evidence above null.

**Figure 7.**
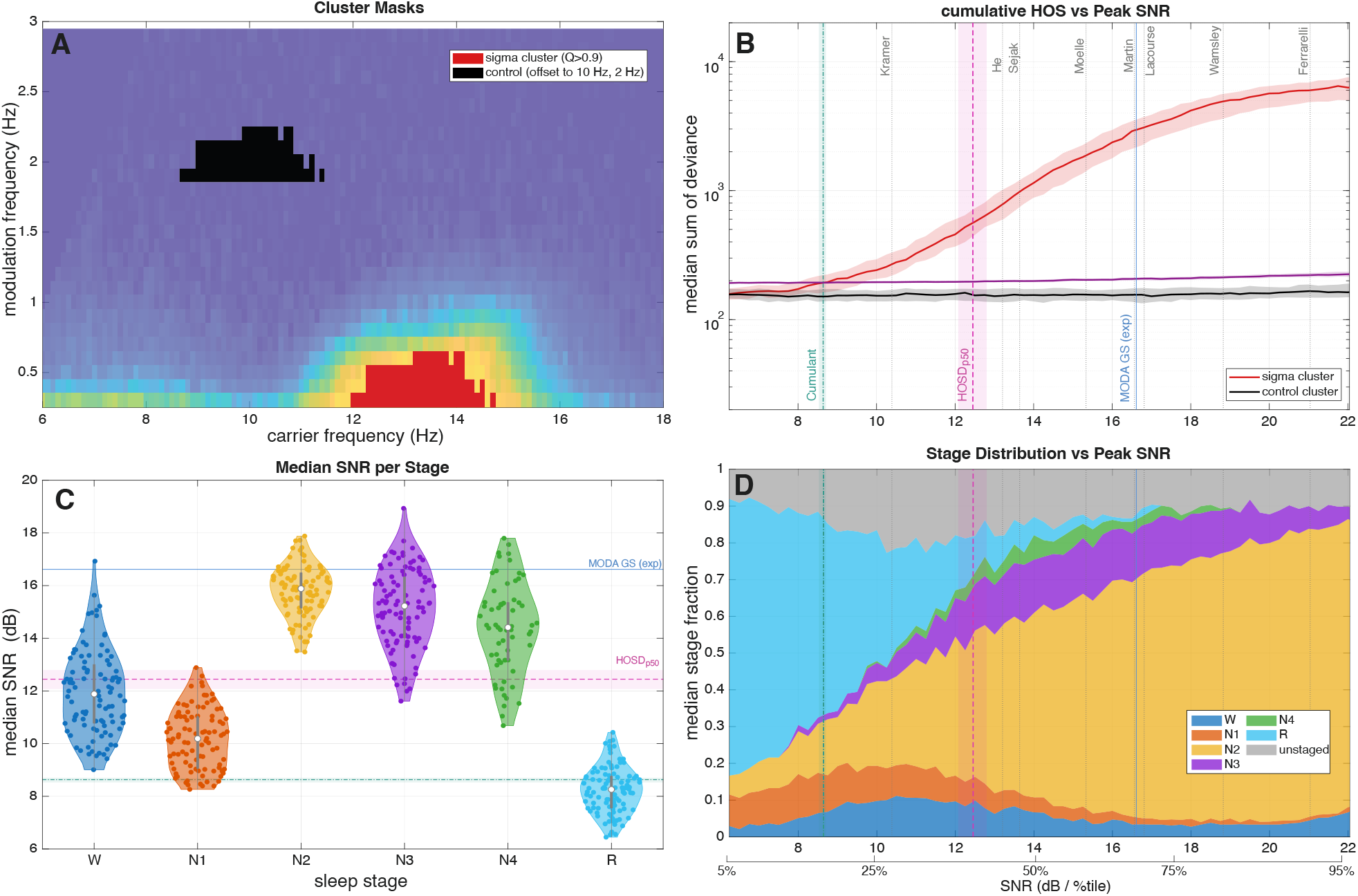
Evidence for spindles below the median detection threshold (“occult” activity) in SNR-sorted data. **A:** For each of the 100 MASS datasets, the sum of deviances for HOS estimates, obtained from the intercept-only complex general linear model, were computed within the region of the modulogram encompassing those coefficients for which at least 90% of datasets exhibited a significant deviation from zero (FDR *q <*= 0.05), which corresponded well to sigma band modulation. A control cluster (*black*) was designated by shifting the mask for the sigma-band cluster to a region with few significant coefficients. **B:** For each dataset, all 10 s analysis windows used in computing the modulogram were sorted according to the maximum SNR normalized amplitude of the filter output within the window. The sum of deviances within the respective clusters was computed for the GLM fit on the subset of sorted windows at or below the given SNR. The median of this value across datasets at each window SNR (*red line*) and inter-quartile range (*shaded*) are plotted with the median *P* = 0.05 significance threshold (*purple line*). Vertical lines indicate the median SNR detection thresholds, calculated as the SNR threshold above which windows contained detections for the given responder, including the cumulant threshold (*teal*); the HOSD_p50_ threshold (*magenta*); MODA GS (*blue*) and other responders are as indicated. The cluster significance threshold was crossed, approximately, within the vicinity of the cumulant threshold, but evidence for spindle-band modulation accumulates well before the HOSD_p50_ window-crossing threshold. **C**: Median peak window SNR broken down by sleep stage. Horizontal lines indicate median thresholds as in B. **D**: Proportion of windows assigned to each stage, computed as a 360-segment moving average of state indicator variables over sorted segments. Low-SNR segments below the cumulant threshold for spindles are dominated by REM, whereas high amplitude windows are dominated by N2. A broad transitional zone is observed between the cumulant-derived threshold and the consensus threshold for expert scorers wherein the dominance of N2 emerges gradually, implying the low-amplitude spindle activity across all stages.

## Discussion

This work establishes the trispectrum as a tool for identifying sleep spindles, regarded as a transiently modulatied carrier oscillation in the sigma band. In this setting, a two dimensional subdomain of the trispectrum conveys information about the presence and spectral characteristics of a modulated oscillation, represented graphically as the modulogram [68; 69]. The spindle-associated peak within the modulogram provides a remarkably stable correlate of spindle activity: it is present in every one of the 108 recordings examined across the DREAMS and MASS databases. In addition to identifying the presence and characteristics of oscillatory bursts, the decompositional approach of HOSD [64] exploits information in the trispectrum to recover a filter matched to the spindle waveform. Finally, a threshold motivated by the relationship of the trispectrum to kurtosis, provides a principled and data-driven criterion for isolating the signal in time, as well as frequency. HOSD-based detection combines three qualities that commonly exist in tension with each other: it renders salient signal properties in an interpretable form; it imposes few necessary assumptions beyond identifiability of HOS; and it isolates a weakly specified target from noise and competing background in a computationally tractable way. These qualities afford a tool that is, to our knowledge, the first to successfully identify sleep spindles in a manner that is fully independent of human judgment.

An important and unique aspect of this method is that it *does not require identifying candidate spindle events as a preliminary step*; rather, spindle waveforms features are recovered from time-averaged HOS before any candidate events are identified in a manner that is not explicitly or implicitly conditioned on any prior notion of what a spindle should look like. Previous accounts of spindle morphology begin with a set of candidate spindles, so that the specificity and biases of the initial detector affect subsequent description of the signal. By reversing the order of identification and detection, the present approach gains two advantages: first, it provides an unbiased view of the aggregate signal, and second, it yields a filter that is optimal for detecting the target signal by being matched to the signal with respect to both phase and amplitude spectra. The present framework also opens numerous additional avenues for extension and development. We develop a Bayesian approach to separating signal and noise through a two-component mixture model, which supplements the permissive kurtosis-based detection criterion with a criterion that allows for detection to be tuned according to a Bayesian decision threshold. The present work therefore complements rather than supplants probabilistic approaches, including hidden Markov models, creating a broad plane for further refinement and development.

Because estimates of HOS suppress Gaussian noise through time-averaging, they recover the spindle waveform in a statistically consistent manner with asymptotically negligible bias. As mentioned, the waveform estimate does not depend on and is not derived from an initial detection of candidate events, which is an important source of bias in prior methods. A noteworthy example of the advantage this provides here is the identification of intra-spindle deceleration as a highly reliable aggregate property of spindles in the MASS database (median −3.9 Hz/s, negative in 98 of 100 MASS recordings; Cohen’s *d* = −2.5), which is visible both in the recovered feature waveform (Fig. 3) and the carrier-dependent group delay seen in the grand average phase modulogram (Fig. 2C). It has previously been noted that spindles tend to exhibit a negative frequency slope [56; 58; 59; 82], including within the MASS database [83; 84]. However, these reports arrive at substantially lower estimates of deceleration than observed in the present analysis: within MASS data, Shimizu et al. (2024) found the per-spindle instantaneous frequency slope to be significantly negative (mean near −0.5 Hz/s), but with a marginal effect size (Cohen’s *d* ≈ −0.3) [83]. Palliyali et al. (2015) similarly found per-dataset mean slopes near −0.7 Hz/s with a moderate effect size over datasets (*d* ≈ −1.6) [84]. This discrepancy likely arises because the per-spindle instantaneous frequency (IF) is an amplitude-weighted average of the target IF and that of concurrent noise [85], which necessarily biases the slope estimate toward zero. Estimating frequency slope through HOSD rather than from individual estimates overcomes this bias. For spindle metrics to have any utility as clinical biomarkers, it is crucial for them to be observed reliably at the individual subject level. Measuring spindle slope with the reliability demonstrated here therefore represents an improvement of kind rather than an increment of degree.

As a method for detecting spindles, the present method performed comparably to non-blind comparison detectors when compared against the MODA expert consensus (Fig. 4A), which is notable in itself. Our introduction of a latent class analysis to the problem, however, adds some nuance to the discussion over which detector represents the “best” in this context. Rather than privileging one responder as the GS over others, the LCM identifies the latent category that best explains the performance of all responders. Comparing LCM-derived performance in Fig. 4C to the expert GS in 4A, it can be observed that responders gravitate under the LCM toward a common performance-recall curve. This suggests that all responders discriminate the same or very similar types of events, with differences between them largely attributed to operating threshold. In this setting, “best” depends on the relative cost of false positives and false negatives in a given application, and there is little basis for declaring one responder the winner without the needed context.

We find that the kurtosis-based threshold employed by HOSD is a permissive one: it reflects the threshold at which evidence of the signal pattern revealed by the trispectrum of the original signal is removed from the trispectrum of the residual. As such, HOSD provides a large pool of candidate events of varying amplitude, after which a suitable Bayesian operating threshold can be identified by fitting a mixture model. HOSD still detected a substantially greater number of events than human scorers and comparison detectors at the Bayesian 0.5 posterior threshold. Although we cannot definitely claim whether the excess detections recovered by HOSD should be regarded as spindles, two separate lines of evidence support the proposition that oscillatory bursts appear below conventional detection thresholds in the spindle band and that, of those, at least a substantial proportion match the AASM spindle definition. First, four sleep specialists evaluated a pool of candidate events generated by HOSD and other detectors in DREAMS excerpts, which also included a high proportion of non-spindle foils to discourage any bias towards affirming detections. Judged against these evaluations, the precision of HOSD was modest, around 0.5, meaning that only about half of the events detected by HOSD at the cumulant threshold survived evaluators’ scrutiny, which was lower than other compared responders. However, recall, at around 0.7, was substantially and significantly higher than that of all other comparison detectors, resulting in the highest *F*_1_ overall, significantly so against every detector except OpenSpindleNet (Fig. 6A). Extra events recovered by HOSD were endorsed by evaluators in approximately 40% of cases, compared with a baseline endorsement of less than 10% for foil events. Events detected by HOSD but missed by all other responders constituted about 36% of the non-foil events endorsed by the evaluators. Precision of HOSD detections may be improved by applying a threshold on event amplitude; yet, with respect to *F*_1_, the decline in recall generally outweighed the rise in precision, so that the *F*_1_-optimal operating point lay at or near the default (cumulant-derived) threshold applied by HOSD. Whether the large number of additional detections should be counted as spindles is however a matter of definition that we do not claim to resolve but that will require further scrutiny by the field.

The second line of evidence for sub-threshold spindle activity is provided by an examination of the point at which cumulative HOS estimates statistically diverge from the null distribution across analysis windows that have been sorted by peak amplitude of the feature detection filter (Fig. 7). This analysis identifies the amplitude at which evidence of the signal identified by HOSD emerges in the modulogram, which may be compared to the detection thresholds of the various responders. Responder thresholds were obtained by finding the amplitude at which at least 50% of windows contain one or more detections from the respective responders. The cluster permutation test statistic used in this analysis, computed cumulatively over sorted analysis windows, crossed the *p* = 0.05 significance threshold of the phase-randomized null distribution at an amplitude that coincided with the cumulant threshold (median 8.66 dB), roughly 4 dB below the natural (*P*_post._ = 0.5) Bayesian threshold and 8 dB below the MODA expert consensus threshold (median 16.6 dB). Moreover, the interval between the cumulant threshold and the MODA expert threshold encompassed approximately 50% of windows; that is, among all sequential 10 s intervals, the detection signal crossed the cumulant threshold in roughly 80% of intervals, while the MODA expert consensus was triggered in roughly 30%. Both results yield compelling evidence that oscillatory bursting in the spindle band appears well below the amplitude at which spindles are conventionally detected and also below the threshold at which a Bayesian operator would confidently identify individual spindle events.

These observations may explain the difficulty of identifying a GS for spindle detection, which has remained an unresolved problem for over a decade [11; 12; 13]. The most commonly accepted GS, expert scoring, is only moderately reliable across scorers (inter-scorer *F*_1_ ≈ 0.61 [10]); a false positive charged to a detector therefore stands a reasonable chance of having in fact been a false negative by the expert. The evidence presented here strongly suggests that the expert-scorer standard is systematically conservative, containing a high proportion of false negatives. By characterizing spindles without reference to any human label and providing a robust human-independent statistical validation of the spindle construct as a genuine signal phenomenon, the present work develops a principled framework for clarifying and addressing the question [11].

The sensitivity of HOSD provides an opportunity to study the relationship between spindle amplitude and sleep stage. Spindles are classically and most prominently associated with N2 sleep, but it is possible that this association has to do with relative amplitude, and thus detectability, rather than density. By stratifying HOSD-derived detections according to amplitude, we obtained a few novel insights into this question, observing that the conventional association with N2 was driven almost entirely by high-amplitude events, above 11 dB. The majority of events identified by HOSD, however, fell bellow this threshold and showed little discrimination between NREM stages and W, while the suppression of spindles in REM sleep appeared at every amplitude (Fig. 5). We conclude that the conventional identification of spindles with N2 is likely to reflect, to a significant degree, an association with spindle amplitude rather than density.

Several limitations deserve to be noted. First, the present approach is based on time-averaged statistics, thus it necessarily provides an aggregate view of spindle characteristics which need not be representative of how individual spindles behave, and which may aggregate over otherwise distinguishable subtypes, such as high and low frequency spindles. While frequency deceleration emerges as a robust property of the spindle waveform in the aggregate, the present analysis provides little insight into the variability of deceleration at the level of individual spindles.

The aggregate estimate, and the application of a single threshold across the recording, both assume stationarity of the signal and background statistics. While we expect some robustness to violations of stationarity in the signal for reasons given in 2.3, robustness to non-stationarity of the background remains an open question. For example, it is possible that the observed global suppression of spindles in REM reflects a global reduction of both signal and noise amplitude during REM rather than a change in oscillatory bursting, as such, although the finding of weak sigma-band modulation during REM in Fig. 2E also tends to support latter conclusion. HOS estimation often also requires generous amounts of data (suggested here to be at least 10 min), so the method is unsuited to short observation windows or strongly non-stationary signals. There are several ways the present analysis might be adapted to better model non-stationarity, including by specifying thresholds that adapt to state, which we leave for future development.

Estimates of HOS are also sensitive to outliers such as large-amplitude transients, so that data must be suitably pre-processed to obtain usable estimates. The present analyses were conducted on a single frontal or central derivation within cohorts of predominantly healthy adults, requiring relatively little pre-processing beyond threshold-based artifact rejection. Although we made every effort not to cherrypick the data to suit the analysis, it is reasonable to question how representative data obtained from a curated open dataset might be of real-world recordings. A signal-agnostic method has clear potential for characterizing atypical spindles in clinical populations, but we present no evidence here on its generalizability beyond the present curated datasets of healthy adults.

It should also be noted that although the method exploits information in higher-order signal statistics to inform filter design and threshold selection, the mechanics of detection remain firmly in the familiar domain of linear filtering and thresholding. A more comprehensive solution to the problem would develop a principled framework for nonlinear filtering, which is conceptually and computationally a more formidable problem than the one addressed here.

Finally, within the cohort of healthy adults observed here, spindle characteristics identified with HOSD are remarkably stable across participants, prompting the reasonable question whether the application of an unsupervised method adds anything of value. Indeed, the LCM analysis suggests a broad equivalence among the various automated detectors and expert annotation, which appear to differ primarily in operating threshold rather than the quality of discrimination or the nature of the event each detects. By identifying a specific and objective signature of spindle-associated oscillatory modulation, the present study contributes independent confirmation of spindles as a distinct electrographic phenomenon and validates existing definitions and methods of identifying them. At the same time, AASM criteria for spindles do not touch on amplitude and so provide no basis on which to arbitrate the question of operating threshold. All detectors considered here are therefore more-or-less equally valid under the existing criteria. The present work highlights operating threshold as a critical source of ambiguity in the definition of sleep spindles and the difficulties that arise from an implicit reliance on a fixed operating threshold, a problem which Purcell et al. (2017) describe as “perhaps, the most pernicious measurement problem” in studying spindles [12]. By circumventing the dependence on detection in measuring spindle characteristics and providing a lower bound on the range of operating thresholds, according to the amplitude at which evidence of spindle-related modulation emerges in the data, HOSD establishes a useful framework for addressing the problem.

## Data availability

The PSG recordings analyzed in this study are openly available: the Montreal Archive of Sleep Studies (MASS) at http://ceams-carsm.ca/mass and the DREAMS Sleep Spindles Database at http://www.tcts.fpms.ac.be/~devuyst/Databases/DatabaseSpindles. Expert spindle annotations from the MODA project are available at https://github.com/klacourse/MODA.

## Code availability

Custom code implementing higher-order spectral decomposition (HOSD) and the analyses reported here is available at https://github.com/ckovach/HOSD. Further scripts are available from the corresponding author on reasonable request.

## Author contributions

C.K.K. conceived the method, performed the analyses and wrote the manuscript. S.V.G., S.K., J.G., O.C., E.W.T., J.A.T., C.A.K., and A.A. advised on analysis and assisted in writing the manuscript. L.C.W., J.L., M.O.S, C.A.K. reviewed PSG data. All authors reviewed and approved the manuscript.

## Funding

This work was supported by grant 3UH3NS113769 from the National Institute of Neurological Disorders and Stroke (NINDS), National Institutes of Health.

## Competing interests

C.K.K. is a named inventor on US Patent 11,159,258 (“Pattern and delay recovery with higher-order spectra”), assigned to the University of Iowa Research Foundation, which relates to the higher-order spectral decomposition method used in this work. The remaining authors declare no competing interests.

## Notes

https://zenodo.org/records/2650142

https://github.com/ckovach/HOSD/

https://borealisdata.ca/dataverse/MASS

