## Supplementary material for "Genuinely Blind Identification of Sleep Spindles through Trispectral Modulation Analysis"

### Spindleness score

Because features identified by HOSD are recovered under weak constraints on frequency and duration, they may not always represent spindles. As a general measure of how well a given feature matches the expected characteristics of a spindle, we computed the following score on the feature waveform weighted by the magnitude spectrum of the detection filter, $\tilde{f}_n = \mathscr{F}^{-1}\{F|H|\}$, where $F$ is the feature's DFT and $H$ is the detection-filter response. Weighting served to suppress residual background noise that survived averaging in the estimated waveform. The score was computed as the harmonic mean of a frequency criterion score, $C_f$, and a duration criterion score, $C_d$, where $C_f$ is the sigma-band-weighted integral of the unit-normalized PSD, $P(\omega) = |\hat{\tilde{f}}(\omega)|^2 / \sum_{\omega'} |\hat{\tilde{f}}(\omega')|^2$, under the following weighting:

$$C_f = \sum_{\omega} b(\omega)\, P(\omega), \quad b(\omega) = \begin{cases} 0 & |\omega| \leq 9 \text{ Hz} \quad \text{or} \quad |\omega| \geq 18 \text{ Hz} \\ 0.5 & |\omega| \in (9, 11 \text{ Hz}) \text{ or } (11, 18 \text{ Hz}) \\ 1 & 11 \leq |\omega| \leq 16 \text{ Hz} \end{cases} \tag{S1}$$

$C_f$ equals 1 if the waveform energy is entirely contained in the conventionally defined sigma band (11–16 Hz); 0.5 if energy lies entirely in one of the 2 Hz transition bands (9–11 or 16–18 Hz); and 0 when no energy falls within 9-18 Hz at all.

For the duration criterion, $C_d$, the normalized waveform estimate was first thresholded using the kurtosis criterion to suppress the effect of any residual background noise in the estimate, and standard width was computed of the thresholded-waveform power, $f_{\text{thr}}$, normalized so that $q(t) = |f_{\text{thr}}(t)|^2 / \sum_{t'} |f_{\text{thr}}(t')|^2$. Width, $W$, was then computed as

$$W = 4\sqrt{\textstyle\sum_n t_n^2 q_n - \left(\sum_n t_n q_n\right)^2}, \quad C_d = \begin{cases} 0 & W \leq 0.25 \\ 0.5 & 0.25 < W < 0.5 \text{ s} \\ 1 & 0.5 \leq W \leq 2 \text{ s} \\ 0.5 & 2 < W < 3 \text{ s} \\ 0 & W \geq 3 \end{cases} \tag{S2}$$

The factor of four sets $W$ to the $\pm 2\sigma$ extent of the envelope, and $C_d$ approximates the AASM duration window for sleep spindles.

The composite spindleness score is taken as the harmonic mean of $C_f$ and $C_d$

$$S = \frac{2\, C_f\, C_d}{C_f + C_d}, \tag{S3}$$

which is zero whenever either criterion is zero: a feature whose duration falls outside (0.25, 3) s, or whose spectral energy falls entirely outside (9, 18) Hz, scores $S = 0$ regardless of how well it satisfies the other criterion. A component scoring partial credit (0.5) on both criteria scores $S = 0.5$, and a component passing both at full weight scores $S = 1$. Within the MASS and DREAMS databases the first HOSD component uniformly achieved a high spindleness score, thus the score was not explicitly applied towards component selection, but is used in the following supplementary analysis to characterize the success of HOSD-based spindle identification under varying quantities of data.

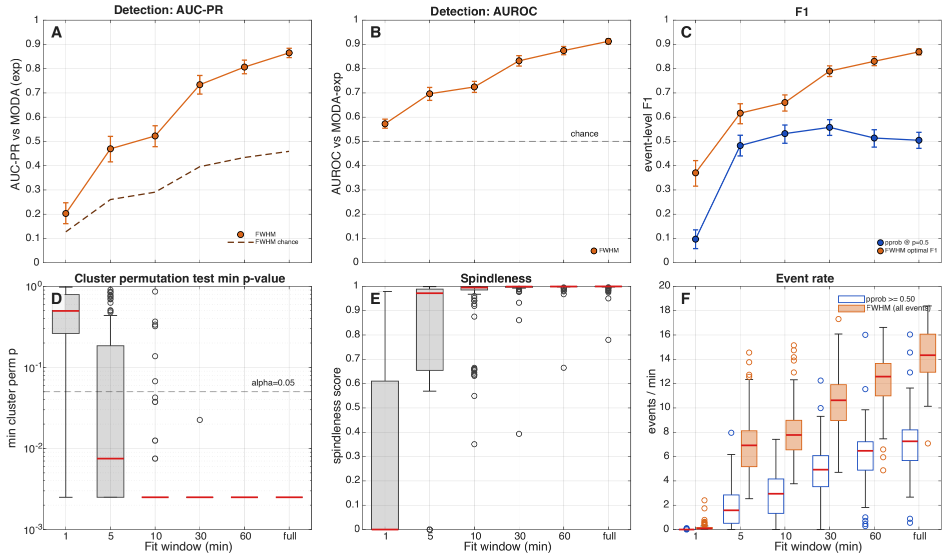


Figure S1: Data quantity needed for HOSD in the MASS data database. For each of 100 recordings from the MASS database, HOSD was fit on windows of varying length 1, 5, 10, 30, 60 min, drawn from a random start position within each subject's concatenated NREM samples (excluding MODA-scored segments), with all available data as an additional comparison. **A:** Event-based area under the precision-recall curve against MODA-expert annotations within the 115-s scored segments, pooled over datasets. Detector events were matched to MODA detections maximum intercept-over-union (IoU) overlap with IoU $> 0.2$. Error bars represent jackknife estimates of population-level s.e.m. computed over datasets. **B:** Event-based AUROC computed in the same way. **C:** Event-based $F_1$ against MODA-expert spindles for two HOSD detectors, $\text{HOSD}_{\text{p50}}$ thresholded at a posterior probability of $p \geq 0.5$ (blue) and $\text{HOSD}_{\text{peak}}$ events at the per-subject optimal $F_1^*$ across the SNR sweep (red). **D:** Minimum cluster-based permutation test $p$-value from the phase-randomized permutation test on the modulogram principal domain (200 permutations; dashed line: $\alpha = 0.05$). Box plots represent the distribution across the 100 datasets (red bar: median; box: inter-quartile range (IQR); whiskers: 1.5×IQR; circles: outliers). **E:** Spindleness score (Eq. S3) of the first HOSD component. **F:** Event rate per minute of the full recording — $\text{HOSD}_{\text{p50}}$ (blue), $\text{HOSD}_{\text{peak}}$ with all detected events (orange).

### Quantity of data needed for reliable detection

The quantity of data required to obtain a reliable estimate of higher-order spectra depends on signal-to-noise ratio but is generally greater than that required for estimates of first and second-order statistics, reflecting both the number of coefficients within HOS and the variance properties of higher moment estimates. To address this question in the present context, we compared performance and quality metrics across a series of durations in the input data, of 1, 5, 10, 30, 60 minutes, and all available data, in each of the 100 MASS sample recordings (Fig. S1). In each case, the input window was drawn with random starting position in the concatenation of N2–N4 NREM samples, excluding the brief MODA-scored segments used for evaluation. The analysis was restricted to NREM to improve the likelihood that sampled segments contained the target signal.

Robustness of the trispectrum estimate was gauged in three ways; first, by examining the frequency with which at least one cluster in the modulogram survived the cluster-based permutation test across all 100 MASS data sets; second, by examining the degree to which the characteristics of the the recovered features those expected of spindles; and third, by examining the agreement of event detections with MODA expert gold standard. The cluster-corrected permutation test reached significance in 8% of data sets at $T = 1$ min, 62% of data sets at $T = 5$ min, 94% at $T = 10$ min, and 100% from $T \geq 30$ min onward.

The recovered feature waveform showed a similarly rapid convergence in its match to the AASM criteria for sleep spindles; the spindleness score (Eq. S3) of the first HOSD component attained a median of $<0.001$ at $T = 1$ min (2% of subjects above 0.95), but rose to 0.97 at $T = 5$ min (60% of subjects above 0.95), 0.995 at $T = 10$ min (83% above 0.95), and 0.999 at $T = 30$ min (97% above 0.95), saturating thereafter, indicating that the recovered feature reliably satisfies the AASM frequency and duration criteria with 10 minutes of N2–N4 NREM data.

Performance-based evaluation suggested a more more graded improvement over all intervals, implying that accuracy of detection continued to benefit from estimates long durations up to the whole recording. Event-level detection performance against MODA-expert annotations continued to rise beyond 10 minutes, with pooled $\text{HOSD}_{\text{peak}}$ AUROC going from 0.57 at $T = 1$ min to 0.70 at $T = 5$ min, 0.73 at $T = 10$ min, 0.83 at $T = 30$ min, 0.88 at $T = 60$ min, and 0.91 at whole-night. Corresponding AUC-PR (Davis & Goadrich, 2006) progressed as 0.20 ($T = 1$ min.) to 0.47 ($T = 5$ min.) to 0.52 ($T = 10$ min.) to 0.73 ($T = 30$ min.) to 0.81 ($T = 60$ min.) to 0.87 ($T =$ whole-night). Pooled $F_1$ for $\text{HOSD}_{\text{peak}}$ at the per-subject F1-optimal SNR threshold rose from 0.37 ($T = 1$ min.) to 0.62 ($T = 5$ min.), 0.66 ($T = 10$ min.), 0.79 ($T = 30$ min,), 0.83 ($T = 60$ min.), and 0.87 ($T =$ whole-night). On the other hand, $\text{HOSD}_{\text{p50}}$ (events thresholded at $p \geq 0.5$) exhibited a more rapid plateau, with pooled $F_1$ rising from 0.10 at $T = 1$ min to 0.48 at $T = 5$ min and $\sim 0.5$ for $T \geq 10$ min (0.53, 0.56, 0.51, 0.51 at $T = 10, 30, 60$ min and whole-night).

These results identify roughly 10 minutes of N2–N4 NREM as a working lower bound for stable HOSD recovery on adult polysomnogram EEG data: cluster-corrected significance is reached in >90% of subjects, the dominant component meets the AASM spindleness criterion in >80%. Detection performance nevertheless continued to improve beyond $T = 30$ min against the MODA expert consensus, suggesting that longer observation windows continue to add value beyond the point when they are adequate for identifying the underlying signal.

### Validity of intra-spindle deceleration estimates

The recovery of spindle characteristics without bias from Gaussian noise is one of the significant advantages of the present method that distinguishes it from available alternatives. To validate this claim for the measurement of spindle deceleration and to test whether the discrepancy between current estimate (-3.9 Hz/s) and those in the literature ($-0.5$ Hz/s [1]; $-0.7$ Hz/s [2]) might be explained by noise-related bias in the latter, we applied the present HOSD pipeline to simulated data containing oscillatory bursts with known deceleration. A one hour sample of data was synthesized at 128 Hz containing simulated spindles in a background of $1/f$ (pink) Gaussian noise. Spindles were modulated by a Gaussian envelope ($\sigma = 0.30$ s, duration $\approx 1.2$ s) with a log-normal amplitude distribution applied to a 13 Hz carrier with a linear chirp of specified slope. The amplitude distribution was selected to match that identified with HOSD in the empirical MASS distribution (median 13.6 dB, IQR 11.7–16.3). The same pipeline was run on the simulated signal as used with the latter and the slope estimated from the recovered feature waveform exactly as in the main analysis. As shown in Fig. S2, the pipeline accurately recovered the true slope across simulated decelerations of $-4$ to $+4$ Hz/s with negligible noise-dependent bias.

For comparison, four previously described methods of slope estimation were applied to the same simulated recordings: (i) the method of Andrillon et al. [3], derived from peri-spindle short-time Fourier spectrograms; (ii) a linear fit the 11–16 Hz analytic (Hilbert) instantaneous frequency over

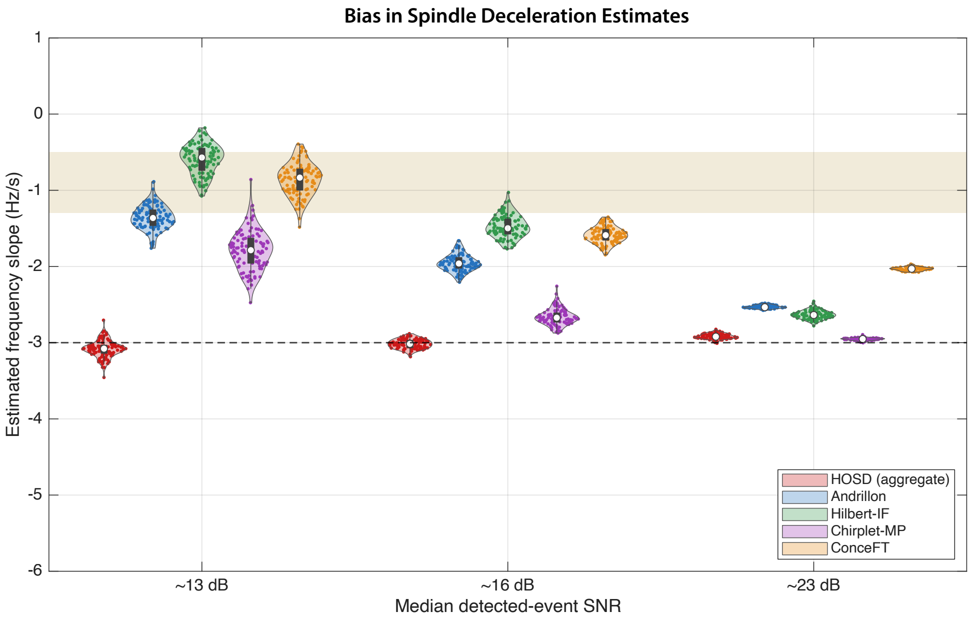


Figure S2: Recovery of spindle deceleration with HOSD. Violin plots show the distribution of frequency slope estimates returned by HOSD and four comparison methods over 100 simulations of one-hour recordings at each of three SNR levels. Recordings contained simulated spindles with −3 Hz/s deceleration (dashed line) in Gaussian noise. HOSD-based estimation (red) closely approximates the true slope at all three SNRs, the first matching the full HOSD detection set (∼13 dB), the second matching the median SNR of MODA expert-detected spindles (∼16 dB), and the third at the upper end of the observed SNR range (∼23 dB). Comparison methods representing of state-of-the-art are: (1) the STFT-based approach of Andrillon et al. (blue), (2) linear fit to the analytic (Hilbert) instantaneous frequency (green), (3) Gabor–chirplet matching pursuit (purple), and (4) ConceFT method of Shimizu et al. (orange). Because these methods all estimate spindle slope by aggregating over measurements on individual events, they are necessarily influenced by the noise background and systematically underestimate slope. Resulting biased estimates are consistent with the range reported in the literature (shaded brown, −0.5 to −1.3 Hz/s), approaching true value only at high SNR.

time [4]; (iii) matching pursuit [5] using a dictionary of chirped Gabor atoms [6, 7, 8]; and (iv) the ConceFT method of Shimizu and Wu [1], which extracts the slope of the instantaneous-frequency ridge from a multitaper time–frequency representation.

All four methods derive the slope estimate from time-frequency decompositions of signal power or analytic instantaneous frequency (IF) obtained from a Hilbert transform, both of which combine the instantaneous frequencies of signal and noise processes weighted by their relative power [4]. Each method therefore produces an estimate that is biased toward zero in the presence of Gaussian noise. The HOSD-derived estimate is minimally biased because the higher-order cumulants of Gaussian noise vanish in expectation and contribute no systematic error to the HOS statistics. As a result, comparison methods grossly underestimated the −3 Hz/s true slope at an SNR chosen to match the median of events identified by HOSD in MASS data (∼13 dB): the median across 100 realizations ranged from −1.8 Hz/s (matching pursuit) to −0.6 Hz/s (IF linear fit), compared to −3.1 Hz/s obtained with HOSD. This deviation exhibited the expected dependence on SNR: at an SNR matching the MODA expert detection threshold ($\sim$16 dB), estimates were still well short of $-3$ Hz/s, ranging from $-2.7$ to $-1.5$ Hz/s, and approached the correct value only at high SNR ($\sim$23 dB), ranging from $-2.95$ to $-2.03$ Hz/s. A method-by-SNR interaction was confirmed with a 2-way ANOVA ($F(8, 1485) = 960$, $p \ll 10^{-6}$, partial $\eta^2 = 0.84$).

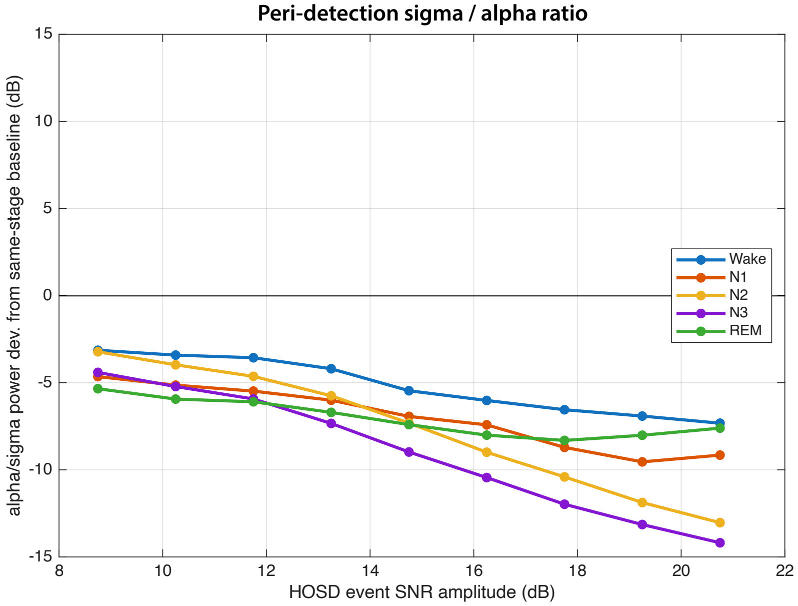


Figure S3: Low amplitude HOSD events are not explained by alpha-band intrusion into the sigma band. For the 697,273 HOSD events across 100 MASS recordings, the deviation, in decibels (dB), of each event's alpha/sigma power ratio (alpha/mu 7–11 Hz, spindle 12–16 Hz) from the median ratio of its own sleep stage in the same recording, plotted against binned event amplitude, separated by stage. Negative values indicate that the ratio of sigma to alpha power is greater than that observed at baseline, consistent with a spindle event; a positive value would suggest that the intrusion of power from oscillations centered in the alpha band resulted in predominantly spurious detections at the given amplitude. Every curve remains negative across the full amplitude range, meaning that median detections are unlikely to be explained by alpha intrusion at any amplitude.

### Validity of low-amplitude detections

A remaining question is whether the low-amplitude events that HOSD recovers might reflect the intrusion of oscillatory activity in adjacent bands; in particular, alpha acivity. To address this question, we examined whether the ratio of power in the alpha and sigma bands deviated from the pattern expected for a sigma-band centered oscillatory burst at any amplitude threshold. For all HOSD events pooled across the 100 MASS recordings, we compared the ratio of a sub-spindle band (alpha, 7–11 Hz) to that in the spindle band (12–16 Hz), normalized to the baseline ratio throughout the corresponding sleep stage. Expressed in dB, a negative deviation from the median indicates relative enhancement of power in the spindle band as might be expected for a genuine spindle, while a positive deviation suggests that intrusion of an alpha oscillation into the sigma band. As shown in Fig. S3, median deviation was negative at all amplitudes and in all stages, supporting a sigma-band specific modulation of power as the predominant source of detections at all amplitudes. One-sample $t$-tests confirmed that the dominance of sigma over alpha power was significant in every amplitude bin (each $p \ll 10^{-6}$; with $t(99) = -25.6$ for the bin with the smallest effect) as well as every stage ($p \ll 10^{-6}$ with $t(99) = -14.0$, for wake in the bin with the smallest effect).
